# L-asparaginase treatment induces a tumor metabolic plasticity and reveals a vulnerability to PARP1/2 inhibitor Olaparib, in B-cell lymphomas

**DOI:** 10.1101/2025.10.24.683536

**Authors:** Aussel Anaïs, Nemazanyy Ivan, Vandenberghe Ashaina, Meola Pauline, Dorat Noémie, Paul Rachel, Habbouche Lama, Exposito Laurie, Nottet Nicolas, Virginie Petrilli, Chiche Edmond, Mounier Nicolas, Marchetti Sandrine, Chaveroux Cédric, Ricci Jean-Ehrland, Chiche Johanna

## Abstract

Intrinsic mechanisms driving secondary resistance to L-asparaginase (ASNase) treatment remain poorly understood. Using a preclinical model of B-cell lymphomas (BCL), sensitive to ASNase, we observed tumor relapse during ASNase treatment despite an initial antineoplastic response. Through *in vivo* metabolic profiling of BCL and stable isotope resolved metabolomics *in vitro* and *in vivo*, we show that ASNase induces a flexible metabolic reprogramming marked by increased *de novo* serine biosynthesis. This adaptive response is driven in part by phosphoglycerate dehydrogenase (PHGDH), which mitigates ASNase-induced reactive oxygen species (ROS) and ROS-mediated DNA damage, enabling tumor cell survival under therapeutic pressure. ASNase-treated malignant B cells exhibit features of replication stress, with activation of the ATR-dependent DNA damage response and increased poly(ADP-ribose) polymerase 1/2 (PARP) activity. Targeting initiation of DNA repair mechanism using the PARP inhibitor Olaparib, enhances the antineoplastic effect of each individual mono-chemotherapy, both *in vitro* and *in vivo.* Importantly, we showed that this therapeutic combination was also effective in homologous recombination (HR)-proficient colorectal cancer cells, supporting its broader therapeutic relevance beyond lymphoid malignancies. Altogether, our findings reveal redox- and DNA repair-dependent metabolic vulnerabilities in ASNase-treated B-cell lymphomas and provide proof-of-concept for a rational combination therapy using clinically approved agents, ASNase and Olaparib.

## Introduction

Cancer cells exhibit enhanced metabolism to meet expanding energetic and biosynthetic demands, a vulnerability that prompted several laboratories to develop metabolic inhibitors for clinical use in oncology ^1–3^. Despite successful evaluations on preclinical models of cancers, only a few of these inhibitors have achieved FDA approval, due in part, to significant toxicities and development of resistance mechanisms ^1,4^. The intrinsic mechanisms by which tumor cells adapt to metabolic targeting *in vivo*, remain poorly understood, thereby limiting the development of additional therapeutic strategies for patients who exhibit resistance to routinely used anti-metabolic agents in the clinic.

L-asparaginase (ASNase) is the only clinically approved drug targeting cancer cell addiction for a particular amino-acid, successfully integrated into multi-agents’ chemotherapy regimens to treat both childhood and adult T-cell and B-cell acute lymphoblastic leukemias (T-ALL, B-ALL) and NK/T-cell lymphomas (NKTCL). By catalyzing the hydrolysis of the non-essential amino acid asparagine in the bloodstream, ASNase triggers starvation and selective apoptosis of asparagine-addicted cancer cells. To date, clinical use of ASNase is restricted to ALL and NKTCL, as these hematological malignancies commonly harbor an epigenetic silencing of the gene encoding the asparagine synthetase that catalyzes the ATP-dependent conversion of L-aspartate (Asp) and L-glutamine (Gln) into L-asparagine (Asn) and L-glutamate (Glu) ^5–7^. Consequently, most ALL and NKTCL are asparagine-auxotrophic, which makes them highly vulnerable to ASNase treatment. Therefore, asparagine synthetase has long been considered a primary determinant of cancer cells’ sensitivity to ASNase treatment ^6^. However, recent studies have shown that ASNase treatment also exhibits anti-tumor efficacy in a variety of other malignancies displaying basal heterogenous asparagine synthetase expression ^8–12^. Moreover, our recent work demonstrated that extracellular asparagine prevents its *de novo* biosynthesis in B-cell lymphomas, regardless of asparagine synthetase expression levels ^9^. These findings challenge the conventional view that asparagine synthetase expression alone is a reliable factor predicting tumor sensitivity to ASNase treatment in asparagine synthetase-expressing malignancies ^9^.

Several mechanisms involved in tumor resurgence during ASNase treatment have been described. Due to its bacterial nature, immunization against ASNase may occur during therapy, leading to neutralization of the enzyme activity. This issue is being addressed through the development of 2^nd^ generation of ASNase formulations with reduced immunogenicity ^13–15^. Additional mechanisms primarily involve the upregulation and/or activation of asparagine synthetase in malignant cells, driven by activation of multiple distinct signaling pathways that converge to an ATF4 (Activating Transcription Factor 4)-dependent transcriptional program promoting cell growth ^10–12^. Nutrient release by stromal cells, which fuels leukemic blasts ^16, 17^ along with specific tumor metabolic adaptations, also account for additional reported mechanisms of resistance to ASNase treatment ^18–20^. However, none have yet led to clinically viable alternative treatments for patients who fail ASNase therapy.

We recently demonstrated the antitumor efficacy of a novel ASNase-based anti-metabolic strategy in patients with refractory/relapsed (R/R) diffuse large B-cell lymphoma (DLBCL) ^8^. Despite complete responses achievement, all patients eventually relapsed during or after completed therapy, suggesting that a subset of malignant cells survived and adapted to circumvent treatment.

Our study aimed to uncover additional cancer cell intrinsic metabolic alterations that contribute to tumor relapse during ASNase treatment, with the intent to reveal novel targetable vulnerabilities. Using a preclinical mouse model of B-cell lymphomas (BCL), we modeled secondary resistance following an initial antineoplastic response to ASNase treatment both *in vitro* and *in vivo*. Through comprehensive metabolomic profiling of BCL, complemented by in-depth analysis of altered metabolic pathways, using the stable isotope tracing in both *in vitro* and *in vivo* settings, we evidenced that ASNase treatment induces a dynamic metabolic reprogramming characterized by increased *de novo* serine biosynthesis in malignant cells. Using pharmacological inhibition, shRNA-mediated silencing, and overexpression of PHGDH—the rate-limiting enzyme of the *de novo* serine biosynthesis—we demonstrated that PHGDH plays a critical role in mediating the tumor’s adaptive response to ASNase. Mechanistically, PHGDH activity mitigates ASNase-induced oxidative stress and related DNA damage, thereby enabling malignant cells to resume proliferation during treatment. Importantly, ASNase-induced DNA damage enhances reliance of malignant B cells on poly(ADP ribose) polymerase 1/2 (PARP) activity, revealing a therapeutic vulnerability to the clinically approved PARP inhibitor, Olaparib, which impairs DNA repair mechanisms.

## Results

### ASNase-sensitive B-cell lymphomas undergo metabolic reprogramming during ASNase therapy both *in vitro* and *in vivo*

Previously, we demonstrated that B-cell lymphomas (BCL) relying on oxidative phosphorylation (OxPhos) metabolism for energy production are sensitive to *E-coli* L-asparaginase (ASNase) therapy, whereas glycolytic-dependent BCL exhibit resistance ^8^. In the present study, we engrafted wild-type C57BL/6 mice with OxPhos-dependent Eµ-*Myc* cells (malignant B cells) isolated from two individual transgenic Eµ-*Myc*^Tg/+^ mice (#506 or #688). Seven days later, mice were treated either with Vehicle or ASNase every 48 hours until the lymphoma reached the ethical endpoint (Figures 1a-b). ASNase treatment showed significant anti-tumor efficacy resulting in delayed BCL development (Figures 1a-b and S1a-b). This represents the initial antineoplastic response to treatment. However, despite continued treatment, all mice eventually developed BCL, a result consistently observed across two independent Eµ-*Myc* clones (#506 or #688) showing equivalent initial sensitivity to ASNase treatment *in vivo*. This secondary response reflects a phase of therapeutic failure, mimicking tumor relapse following an initial favorable response, as observed in patients with R/R DLBCL treated with innovative ASNase-based anti-metabolic therapy ^8^.

**Figure 1:**
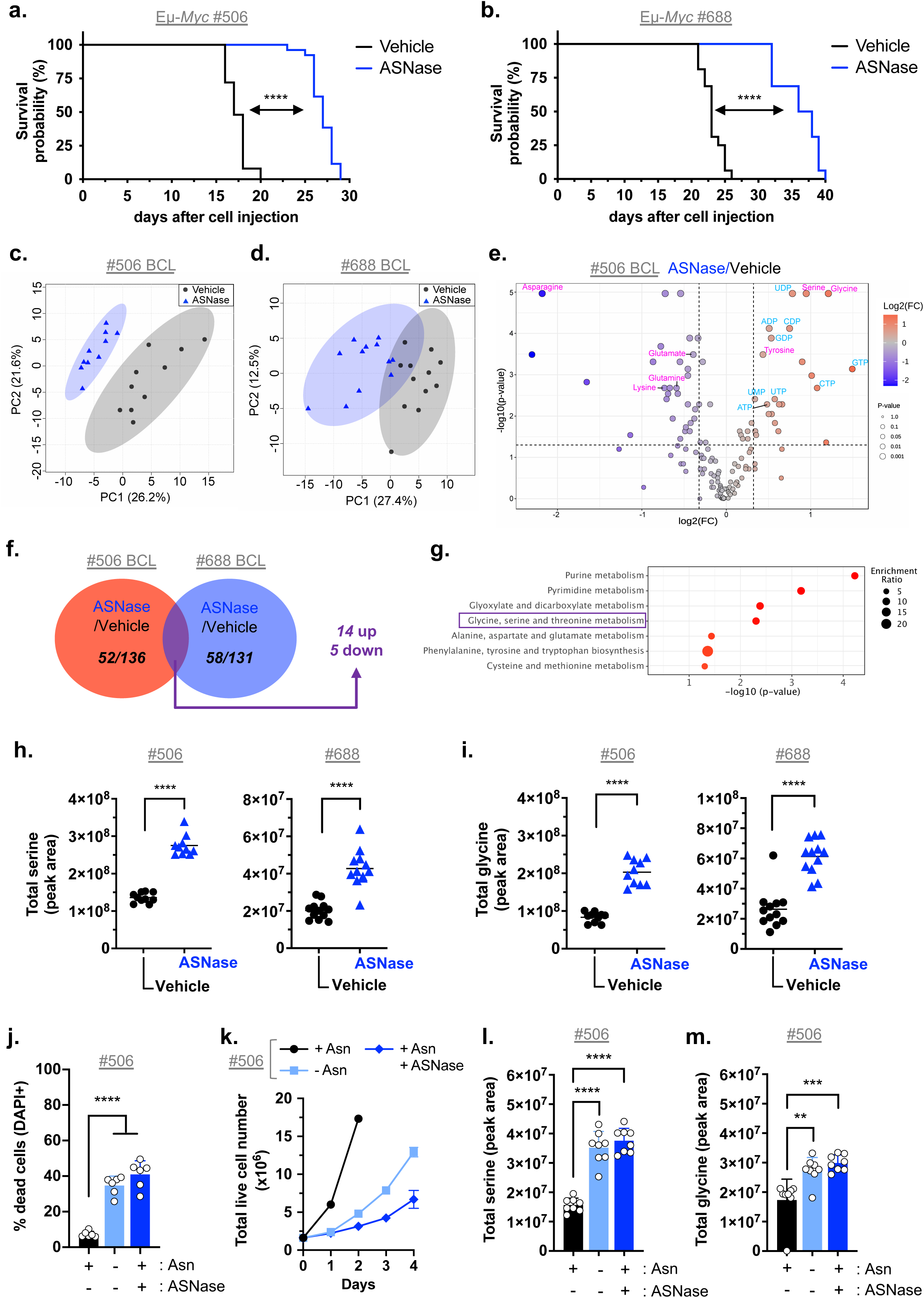
ASNase treatment induces a tumor-associated metabolic response, characterized by increased intracellular serine and glycine levels, both *in vitro* and *vivo*. **a.** Survival curves of syngeneic wild-type C57BL/6 mice bearing OxPhos-dependent Eμ-*Myc* #506 lymphoma treated with Vehicle (NaCl 0.9%) or *E-Coli* L-asparaginase (ASNase), every 48 hours until B-cell lymphomas (BCL) reached the ethical endpoint (n=25 Vehicle-treated mice; n=26 ASNase-treated mice; 3 experiments). P-value from log-rank test. **b.** As in **a.** with primary OxPhos-dependent Eµ-*Myc* #688 lymphoma (n=16 mice/group; n= 2 experiments). P-value from log-rank test. **c.** Principal-component analysis (PCA) of metabolites abundance in axillary BCL harvested C57BL/6 mice bearing Eµ-*Myc* #506 lymphoma, treated with Vehicle or ASNase until endpoint, based on 136 metabolites detected by LC/MS (n=10 mice/group). **d.** As in **c.** from C57BL/6 mice bearing Eµ-*Myc* #688 lymphoma, based on 131 metabolites detected by LC/MS (n=12 mice/group). **e.** Volcano plot highlighting significantly deregulated metabolites (Fold change (ASNase/Vehicle) >1.25 and raw P-value < 0.05) in Eµ-*Myc* #506 cells-derived BCL described in **c.** (n=10 mice/group). Nucleotide precursors labelled in blue, proteinogenic amino acid labelled in pink. **f.** Venn diagram illustrating the number of statistically significant deregulated metabolites obtained from targeted metabolomic analysis of BCL presented in **c.** and **d.** Significance was defined raw P-value < 0.05 and fold change threshold was (ASNase/Vehicle) >1.25. **g.** Overview of top statistically significant enriched KEGG metabolite sets from 19 commonly deregulated metabolites in BCL treated with ASNase *in vivo*, highlighted in **f.** (see Table S1). **h. i.** Relative abundance (peak area) of total intracellular serine (**h.**) and glycine (**i.**) in BCL presented in **c.** and in **d.** (from Eµ-*Myc* #506 cells, n=10 mice/group; from Eµ-*Myc* #688 cells, n=12 mice/group). P-value from t-test. **j.** Percentage of dead Eμ-*Myc* (#506) cells (DAPI+) cultivated for 24 hours in Asn-free medium supplemented with (+) or without (-) Asn (0.37 mM) and ASNase (0.003 IU/ml). Data are expressed as mean ± SD (n= 6 experiments). P-value from t-test. **k.** Proliferation of Eµ-*Myc* (#506) cells cultivated as in **j.** until 96 hours. Data are expressed as the mean number of live cells (DAPI-) ± SEM (n= 6 experiments). **l. m.** Relative abundance (peak area) of total intracellular serine (**l.**) and glycine (**m.**) in live Eµ-*Myc* (#506) cells cultivated for 24 hours under conditions described in **j.** Data are expressed as mean ± SD (n=4 biological replicates and two technical replicates). P-value from t-test. **, p < 0.01; ***, p < 0.001; ****, p < 0.0001.

One mechanism of tumor resurgence during ASNase treatment in clinical settings is the production of anti-ASNase antibodies which neutralize the enzyme and are often associated with allergic episodes in patients. To rule out decreased ASNase activity as the cause of tumor relapse, mice received a final bolus of Vehicle or ASNase at endpoint. Four hours post administration, BCL-bearing mice treated with ASNase exhibited undetectable plasma asparagine concentration and elevated plasma aspartate concentration compared to Vehicle-treated controls, indicating sustained ASNase activity despite disease relapse (Figure S1c). Regardless of the tumor entity or energetic dependency, therapy-resistant cancer cells frequently enhance mitochondrial energetic functions to support survival ^21–24^. Consistently, we observed a significant increase in the contribution of OxPhos to ATP production in BCL that progressed during ASNase treatment *in vivo*, suggesting a metabolic reprogramming to meet elevated ATP demands (Figure S1d).

We next conducted targeted metabolomic analysis in BCL undergoing Vehicle or ASNase treatment *in vivo*. Principal component analysis (PCA) and Venn diagrams showed that the metabolic profile of BCL treated with ASNase *in vivo* was distinct from that of Vehicle-treated BCL, irrespective of the Eµ-*Myc* clone used (Figures 1c-d-e and S1e). Nineteen (14%) significantly deregulated metabolites were common to both datasets (#506-derived or #688-derived BCL) and exhibited similar directional regulation in response to ASNase treatment with 14 metabolites increased and 5 metabolites decreased (Figure 1f and Table S1). These shared metabolites likely reflect core metabolic response to ASNase therapy, that are independent of the Eµ-*Myc* clone origin, suggesting conserved metabolic vulnerabilities in our *in vivo* model. Metabolite Set Enrichment Analysis (MSEA) using the Kyoto Encyclopedia of Genes and Genomes (KEGG) pathway computational tool, revealed deregulated nucleotide and amino acid (AA) metabolism in ASNase-treated BCL (Figure 1g). Among proteinogenic AA, serine, glycine, tyrosine, and asparagine were the only significantly and consistently deregulated AA in ASNase-treated BCL, meeting the fold change threshold (ASNase/Vehicle) >1.25 and raw P-value < 0.05, across both datasets (Figures 1e, S1e and Table S2). Indeed, BCL treated with ASNase *in vivo* exhibit a two-to three-fold increase in steady-state serine and glycine levels in ASNase-treated BCL compared to Vehicle-treated BCL (Figures 1h-i).

To confirm that ASNase treatment triggered serine metabolism remodeling in malignant cells, Eµ-*Myc* cells were cultured in DMEM medium lacking asparagine but containing supraphysiological concentrations of serine and glycine, with (+) or without (-) supplementation of asparagine and ASNase. A concentration of 0.003 IU/ml ASNase was used to avoid hydrolysis of glutamine into L-glutamate (Figure S1f). ASNase treatment significantly increased the death of Eµ-*Myc* cells in 24 hours (Figures 1j and S1g). However, Eµ-*Myc* cells surviving this nutritional stress resumed proliferation over time (Figures 1k et S1h), a process accompanied by a significant increase in intracellular serine and glycine levels following 24 hours of ASNase treatment (Figures 1l-m and S1i-j). To specifically evaluate metabolic adaptations to asparagine deprivation— since ASNase-mediated release of aspartate and ammonium can each influence the tumor metabolic response —we cultured Eµ-*Myc* cells in asparagine-free medium. This experimental setup ensured that the observed metabolic changes were directly attributable to asparagine depletion, thereby reinforcing the robustness of our conclusions. Our findings indicate that ASNase treatment induces a metabolic response in BCL, both *in vitro* and *in vivo*, characterized by elevated serine and glycine levels in malignant B cells.

### Increased *de novo* serine biosynthesis in ASNase-treated B-cell lymphomas, *in vitro* and *in vivo*

Increased total serine levels in malignant cells may result from either enhanced serine import via active transport or increased *de novo* serine biosynthesis from the glycolytic intermediate 3-phosphoglycerate and glutamine-derived glutamate through a 3-step enzymatic process catalyzed by phosphoglycerate dehydrogenase (PHGDH), phosphoserine aminotransferase-1 (PSAT1), and phosphoserine phosphatase (PSPH). Eµ-*Myc* cells proliferation was unaffected by extracellular L-serine/glycine withdrawal, regardless of asparagine availability (Figure S2a). In line with a previous report describing asparagine as an amino acid exchange factor facilitating serine import ^25^, asparagine-restricted Eµ-*Myc* cells exhibited significantly reduced extracellular serine uptake, despite an increase in total intracellular serine levels (Figures 2a and S1i). Moreover, incubation of Eµ-*Myc* cells in serine/glycine-free medium, still resulted in a marked and significant increase in intracellular serine and glycine abundance in ASNase-treated cells compared to control cells, suggesting activation of the *de novo* serine and glycine biosynthesis pathway upon treatment (Figures 2b-c). Supporting this hypothesis, asparagine-deprived Eµ-*Myc* cells upregulated PHGDH, PSAT1, PSPH mRNA and protein expression, along with the expected induction of asparagine synthetase (Figures 2d-e and S2b). The glutamine synthase (GLUL) that has been associated with resistance of sarcoma cells to ASNase ^18, 19^, was upregulated only in glutamine-deprived, but not in asparagine-deprived Eµ-*Myc* cells, suggesting a weak contribution of GLUL to secondary resistance to ASNase treatment in malignant B cells. The expression level of the glutamate oxaloacetate transaminase 1 (GOT1), which remained unchanged upon nutritional stress, indicates a selective regulation of specific metabolic enzymes consistent with upregulation of the transcription factor ATF4 (Figures 2d-e and S2c), as already described ^26–29^.

**Figure 2:**
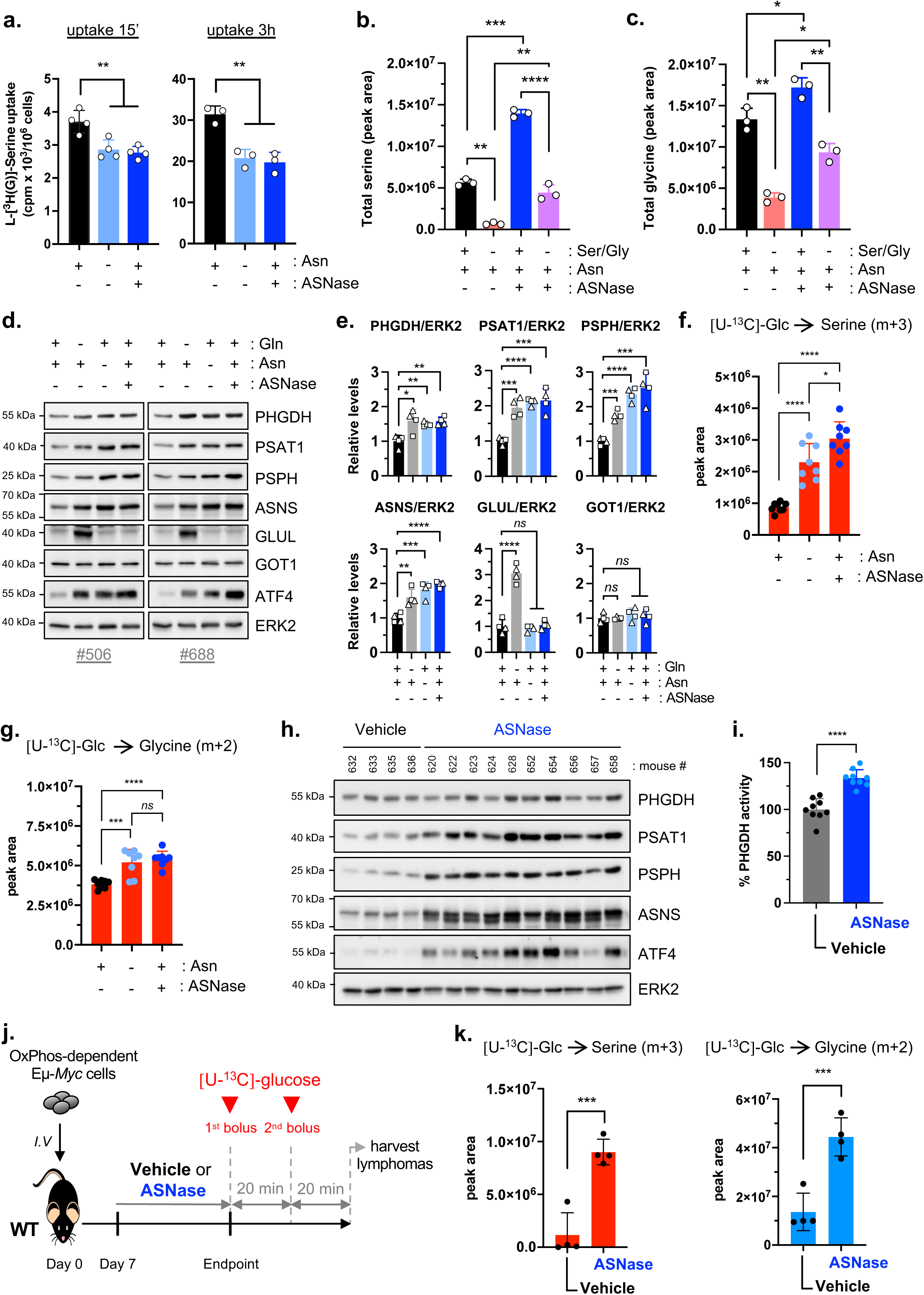
*De novo* serine biosynthesis increased in B-cell lymphomas treated with ASNase *in vivo*. **a.** L-[^3^H(G)]-serine transport rate (cpm/10^6^ cells, resulting from 15 min uptake, left panel or 3 hours uptake, right panel) in Eµ-*Myc* (#688) cells following 24 hours incubation in serine and glycine-containing medium, with (+) or without (-) Asn (0.37 mM) and ASNase (0.003 IU/ml). Data are expressed as mean ± SD (n≥ 3 experiments). P-value from t-test. **b. c.** Relative abundance (peak area) of total intracellular serine (**b.**) and glycine (**c.**) levels in Eµ*-Myc* (#688) cells cultivated for 24 hours in Asn, L-serine and glycine-free medium, (+) or not (-) with serine, glycine (Ser/Gly; 0.4 mM/0.4 mM), Asn (0.37 mM) and ASNase (0.003 IU/ml) (n=3 biological replicates). P-value from 2-way Anova followed by Tukey’s test. **d.** Total protein extracts prepared from whole live Eµ-*Myc* #506 (left panel) and #688 (right panel) cells cultivated for 24 hours in glutamine (Gln) and Asn-free medium supplemented (+) or not (-) with Gln (2 mM), Asn (0.37 mM) or ASNase (0.003 IU/ml), were immunoblotted for the indicated proteins. ERK2, loading control. Immunoblots shown are representative of 2 independent experiments for each Eµ-*Myc* line. **e.** Relative quantification of the expression levels of indicated proteins in Eµ-*Myc* #506 (square, n=2 independent experiments) and in Eµ-*Myc* #688 (triangle, n=2 independent experiments) cells, following 24 hours of incubation as in **d.** Data are represented as mean ± SD. P-value from P-value from t-test. **f. g.** Relative abundance (peak area) of ^13^C-labelled serine m+3 (**f.**) and ^13^C-labelled glycine m+2 (**g.**) isotopologues in Eµ-*Myc* (#688) cells cultivated for 24 hours in glucose and asparagine-free medium supplemented with 25 mM [U-^13^C]-glucose and with (+) or without (-) Asn (0.37 mM) and ASNase (0.003 IU/ml). Data are expressed as mean ± SD (n=4 biological replicates and two technical replicates). P-value from t-test. **h.** Total protein extracts prepared from BCL harvested in C57BL/6 mice bearing Eµ-*Myc* (#506) lymphoma, treated with Vehicle or ASNase every 48 hours till disease endpoint, were immunoblotted for the indicated proteins (Vehicle, n=4 mice; ASNase, n=10 mice). ERK2, loading control. **i.** Percentage of PHGDH activity in BCL harvested from Eµ-*Myc* cells-bearing C57BL/6 mice treated as in **g.** Data are expressed as mean ± SD (n=9 mice/group). P-value from t-test. **j.** Schematic representation of *in vivo* [U-^13^C]-glucose consecutive bolus protocol following Vehicle or ASNase treatment in C57BL/6 mice bearing OxPhos-dependent Eµ-*Myc* #688 BCL. **k.** Relative abundance (peak area) of ^13^C-labelled serine (m+3) and glycine (m+2) isotopologues in BCL harvested in C57BL/6 mice bearing Eµ-*Myc* #688 lymphoma, treated as in **j**. Data are expressed as mean ± SD (n=4 mice/group). P-value from t-test. PHGDH, phosphoglycerate dehydrogenase; PSAT1, phosphoserine aminotransferase 1; PSPH, phosphoserine phosphatase; ASNS, asparagine synthetase; GLUL, glutamine synthetase, GOT1, Glutamate-Oxaloacetate Transaminase-1; ATF4, Activating Transcription Factor 4. *ns*, not significant; *, p < 0.05; **, p < 0.01; ***, p < 0.001; ****, p < 0.0001.

Given that the expression level of metabolic enzyme does not necessarily reflect its activity^9^, we assessed the activity of PHGDH—the rate-limiting enzyme in *de novo* serine biosynthesis— as well as the level of *de nov*o-synthesized serine and glycine by tracing [U-^13^C]-glucose in Eµ-*Myc* cells *in vitro*. We observed a significant increase in PHGDH activity (Figure S2d), along with elevated levels of *de novo* ^13^C-labelled serine (m+3) and ^13^C-labelled glycine (m+2) derived from [U-^13^C]-glucose, in asparagine-deprived Eµ-*Myc* cells, even in medium containing supraphysiological concentrations of both amino acids (Figures 2f-g and S2e). These cells also exhibited increased levels of ^13^C-labelled asparagine (m+2 and m+3), consistent with active asparagine synthetase under conditions of extracellular asparagine withdrawal (Figure S2f). Similarly, using α-^15^N-L-glutamine, we detected increased levels of ^15^N-labelled serine (m+1) and ^15^N-labelled asparagine (m+1), further supporting increased *de novo* serine biosynthesis in malignant B cells exposed to an asparagine-free environment (Figures S2e-g-h).

*In vivo*, BCL harvested from ASNase-treated mice also exhibit upregulated PSAT1, PSPH, ASNS and ATF4 expression, though PHGDH expression was not (Figures 2h and S2i). Nevertheless, PHGDH activity was significantly increased in BCL of ASNase-treated mice (Figure 2i). Following several weeks of Vehicle or ASNase treatment, BCL-bearing mice received two consecutive boluses of [U-^13^C]-glucose at a 20 min interval to quantify de novo-synthesized serine, glycine, and asparagine in tumors (Figure 2j). Equivalent enrichment of fully carbon-labeled glucose (m+6) was detected in the plasma of both Vehicle and ASNase-treated mice bearing BCL (Figure S3a). Consistent with *in vitro* data, levels of carbon-labelled ^13^C-labelled serine (m+3), glycine (m+2) and asparagine (m+2) were significantly increased in ASNase-treated BCL, accompanied by elevated total levels of serine, glycine and a corresponding decrease in total levels of asparagine, suggesting increased *de novo* serine/glycine and asparagine biosynthesis from [U-^13^C]-Glucose in BCL exposed to ASNase treatment *in vivo* (Figures 2k and S3-b-c-d-e). As previously reported in WT mice and in multiple murine cancer models ^30^, we also confirmed elevated plasma serine concentration in BCL-bearing C57BL/6 mice treated with ASNase (Figure S3f).

Altogether, our results demonstrate that malignant B cells exposed to an asparagine-restricted environment support *de novo* serine biosynthesis, despite available environmental serine and glycine.

### PHGDH supports ASNase-sensitive malignant B cells proliferation during therapy

We next investigated whether ASNase-sensitive malignant B cells increase *de novo* serine biosynthesis to facilitate their outgrowth in an asparagine-deprived environment. First, we tested several commercially available PHGDH inhibitors—including three allosteric compounds (NCT-503, CBR-5884 and PKUMDL-WQ2101) and one competitive inhibitor (BI-4916) ^31–34^ in Eµ-*Myc* cells to evaluate their effects on PHGDH enzymatic activity and cellular viability. While all compounds exhibited concentration-dependent cytotoxicity *in vitro* (Figures S4a-b-c-d), only BI-4916, a cell-permeable and selective NAD⁺/NADH-competitive inhibitor of PHGDH ^34^, effectively inhibited PHGDH activity in Eµ-*Myc* cells (Figure S4e). Based on these results, BI-4916 was selected for further investigations. Notably, our findings are consistent with prior studies demonstrating the superior efficacy of BI-4916 in inhibiting *de novo* serine biosynthesis in the breast cancer cell line MDA-MB-468, compared to allosteric inhibitors ^35^. Treatment with 10 µM of BI-4916 inhibited PHGDH activity by 80%, significantly reduced total intracellular serine levels (Figures 3a and S4e-f) and induced an average significant 32% increase in Eµ-*Myc* cell death within 24 hours (Figures 3b and S4d-g). When combined with ASNase, BI-4916 further enhanced cell death in a caspase-dependent manner within 24 hours, compared to either treatment alone (Figures 3b, S4g and S4h). Moreover, ASNase/BI-4916 co-treatment significantly delayed the outgrowth of Eµ-*Myc* cells, underscoring the contribution of PHGDH activity to malignant B cells’ adaptation during ASNase treatment (Figure 3c). Similar results were obtained with independent ASNase-sensitive Eµ-*Myc* cells (Figures S4g-i) and with the NAD^+^/NADH competitive PHGDH inhibitor BI-4924, that was chemically modified to generate the esterified prodrug form, BI-4916 ^34^ (Figures S4j-k).

**Figure 3:**
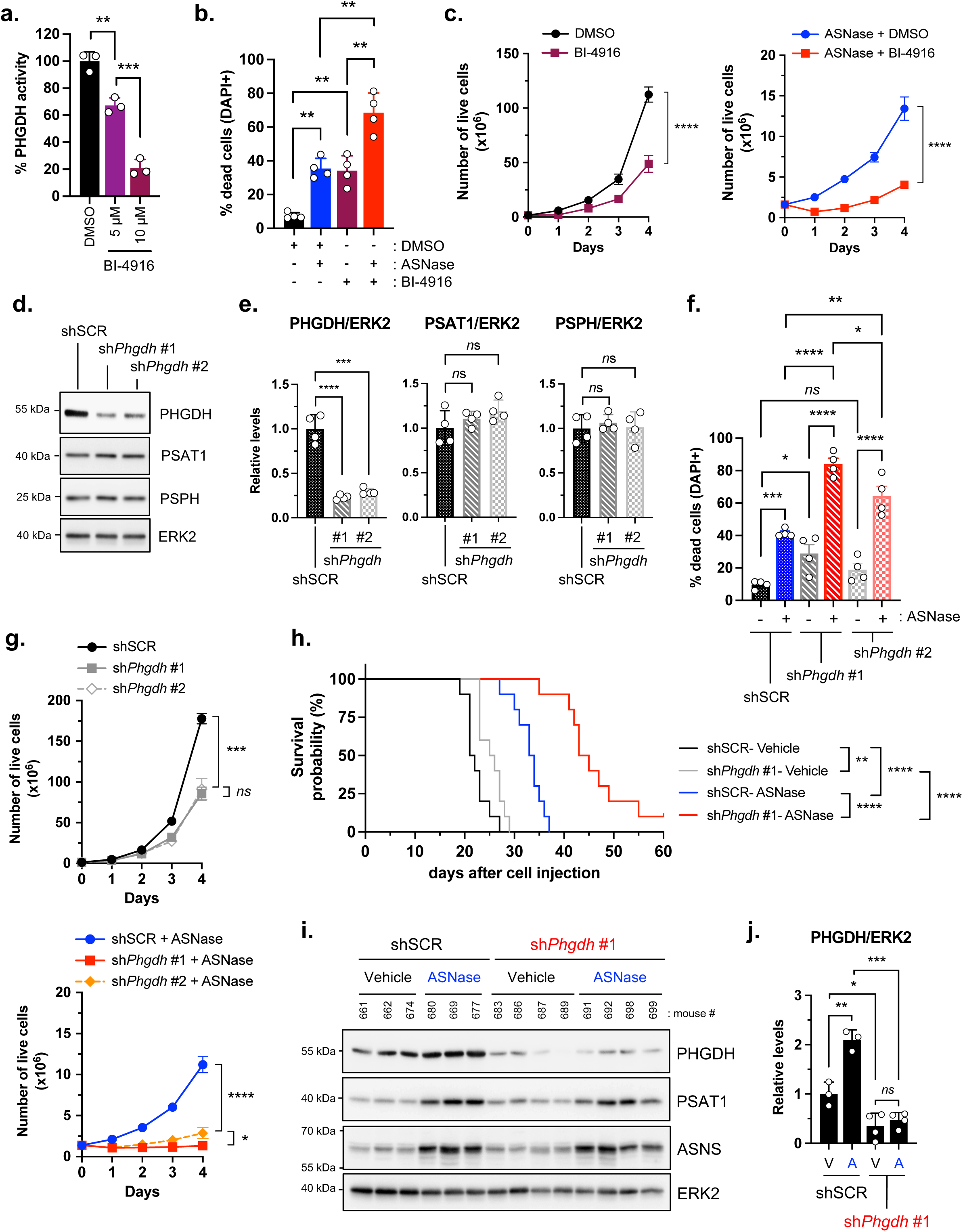
PHGDH contributes to malignant B cell adaptation to ASNase treatment *in vitro* and *in vivo*. **a.** Percentage of PHGDH activity in Eµ-*Myc* (#688) cells treated with DMSO or indicated concentrations of the PHGDH inhibitor, BI-4916 for 24 hours in Asn-containing medium. Data are expressed as mean ± SD (n= 3 experiments). P-value from t-test. **b.** Percentage of dead Eµ-*Myc* (#688) cells (DAPI+) following 24 hours of treatment with DMSO, ASNase (0.003 IU/ml) and/or BI-4916 (10 µM) in Asn-containing medium. Data are expressed as mean ± SD (n= 4 experiments). P-value from 2way Anova, followed by Tukey’s test. **c.** Four-days proliferation of Eµ-*Myc* (#688) cells treated as described in **b**. Data are expressed as the mean number of live cells (DAPI-) ± SEM (n= 4 experiments). P-value from t-test. **d.** Total protein extracts prepared from whole Eµ-*Myc* (#506) cells stably expressing shRNA targeting the firefly luciferase (shSCR) or the murine *Phgdh* mRNA (two independent sh*Phgdh* #1 and #2) were immunoblotted for indicated proteins. ERK2, loading control. Immunoblots shown are representative of 4 independent experiments. **e.** Relative quantification of results presented in **d.** Data are represented as mean ± SD (n=4 independent experiments). P-value from t-test. **f.** Percentage of dead Eµ-*Myc* (#506) cells (DAPI+) silenced (sh*Phgdh* #1 or #2) or not (shSCR) for *Phgdh* mRNA following 24 hours incubation in Asn-containing medium supplemented (+) or not (-) with ASNase (0.003 IU/ml). Data are expressed as mean ± SD (n= 4 experiments). P-value from 2way Anova, followed by Tukey’s test. **g.** Four-days proliferation of Eµ-*Myc* cells silenced or not (shSCR) for *Phgdh* mRNA, treated as in **f.** Data are expressed as the mean of proliferation index ± SEM (n=5 experiments). P-value from t-test. **h.** Survival curves of WT C57BL/6 mice intravenously injected with Eµ-*Myc* (#506) cells stably expressing control shRNA (shSCR) or shRNA targeting the murine *Phgdh* mRNA (sh*Phgdh* #1) and treated 7 days later with Vehicle (NaCl 0.9%) or ASNase every 48 hours until endpoint (n=10 mice/group). P-value from log-rank test. **i.** Total protein extracts prepared from Eµ-*Myc* cells isolated from BCL of C57BL/6 mice intravenously injected with Eµ-*Myc* (#506) cells stably expressing control shRNA (shSCR) or shRNA targeting the murine *Phgdh* mRNA (sh*Phgdh* #1) and treated with Vehicle or *E-Coli* L-asparaginase (ASNase) every 48 hours till endpoint, were immunoblotted for the indicated proteins (shSCR-Vehicle, n=3 mice; shSCR-ASNase, n=3 mice; sh*Phgdh* #1-Vehicle, n=4 mice; sh*Phgdh* #1-ASNase, n=4 mice). ERK2, loading control. V: Vehicle; A: ASNase. **j.** Relative quantification of PHGDH protein levels presented in **i.** Data are normalized to the control condition shSCR-Vehicle and expressed as mean ± SD. P-value from 2way Anova, followed by Tukey’s test. *ns*, not significant; *, p < 0.05; **, p < 0.01; ***, p < 0.001; ****, p < 0.0001.

Due to the intrinsic instability of BI-4924 and BI-4916, which precludes their use in *in vivo* studies ^34^, we sought to confirm that the observed effects were attributable to on-target PHGDH inhibition by stably expressing shRNA targeting the murine *Phgdh* mRNA in Eµ-*Myc* cells. Two shRNA constructs (#1 and #2) effectively reduced PHGDH protein levels without altering PSAT1 or PSPH expression levels (Figures 3d-e). Consistent with pharmacological inhibition, silencing of *Phgdh* sensitized Eµ-*Myc* cells to ASNase-induced cell death and prevented the expansion of surviving cells *in vitro* (Figures 3f-g). *In vivo*, ASNase-treated mice engrafted with Eµ-*Myc*-sh*Phgdh* #1 cells displayed significantly delayed lymphoma progression compared to those bearing control Eµ-*Myc*-shSCR cells (Figure 3h), confirming PHGDH as a key mediator of BCL relapse during ASNase therapy. Of note, at endpoint, we confirmed downregulated levels of PHGDH protein in BCL of Vehicle and ASNase treated mice injected with Eµ-*Myc*-sh*Phgdh* #1 cells (Figure 3i). On the contrary, PHGDH overexpression in Eµ-*Myc* cells accelerated BCL development upon ASNase treatment (Figures S4l-m-n), further validating PHGDH’s role in the adaptive resistance mechanisms.

Collectively, our results highlight the critical role of PHGDH activity and *de novo* serine biosynthesis in the proliferation of malignant B cells exposed to an asparagine-deprived environment.

### ASNase treatment induces a reversible tumor metabolic response *in vivo*

Prolonged nutritional stress, such as that imposed by weeks of treatments with the clinically used ASNase, raises questions about the persistence of mechanisms driving tumor relapse. To assess the reversibility of this adaptation, Eµ-*Myc* cells that survived 4 days of ASNase treatment *in vitro* (1^st^ challenge), were released from therapy (C1 cells). Upon re-exposure to ASNase (2^nd^ challenge), C1 cells displayed an ASNase sensitivity comparable to chemo-naïve parental (WT) Eµ-*Myc* cells, suggesting that the adaptive response is transient (Figures 4a-b-c). This result was consistent across other independent ASNase-sensitive Eµ-*Myc* cells (Figures S5a-b).

**Figure 4:**
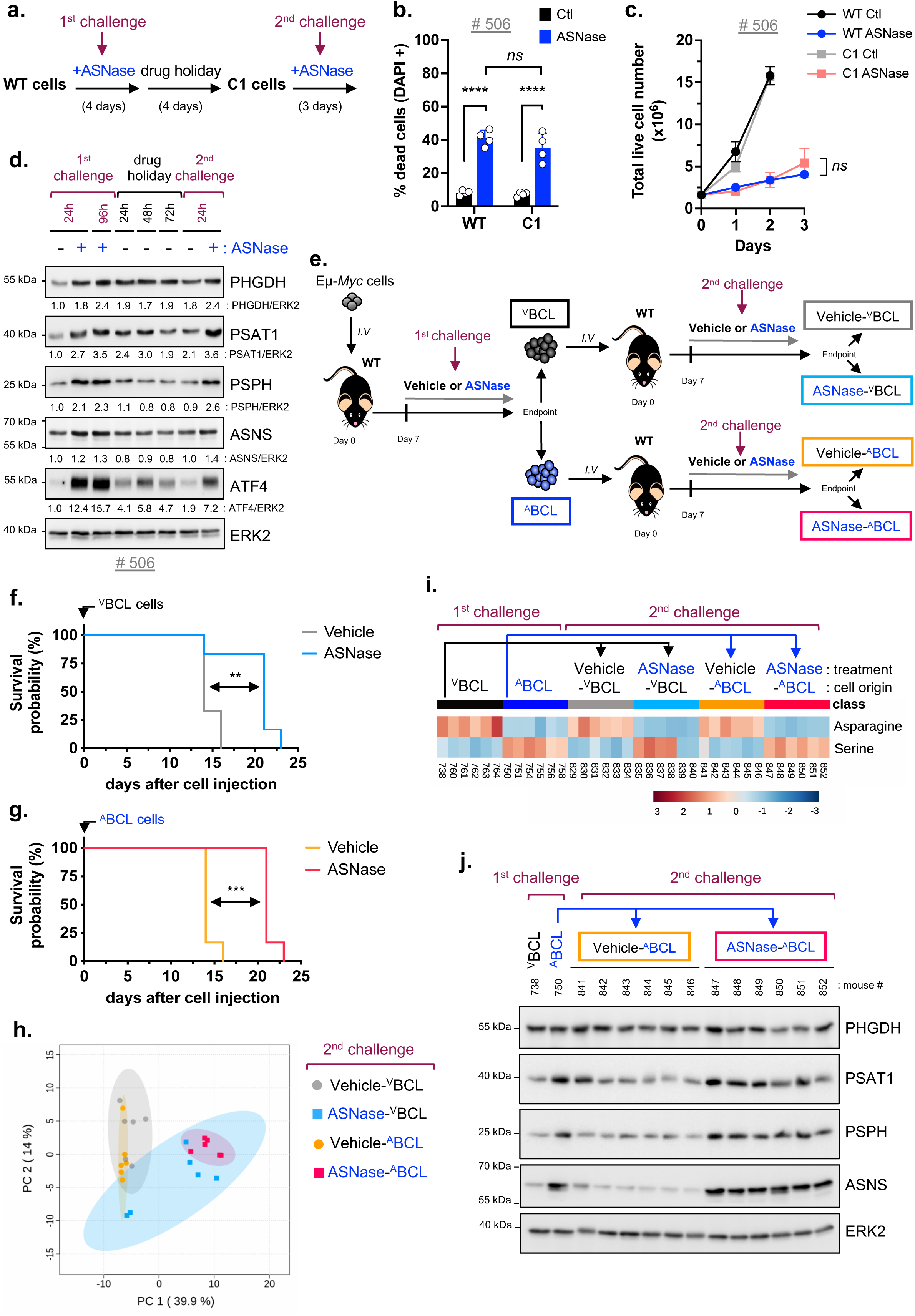
ASNase treatment triggers a flexible tumor metabolic program *in vivo*. **a.** Schematic representation of the experimental design. Following 4 days of ASNase (0.003 IU/ml; 1^st^ challenge), wild-type (WT) Eµ-*Myc* (#506) cells were washed and re-seeded in Asn-containing medium (drug holiday). The resulting cell population, termed C1 cells, was treated with ASNase for 4 days (2^nd^ challenge). **b.** Percentage of dead (DAPI+) WT and C1 cells (from Eµ-*Myc* #506 cells) incubated in Asn-containing medium with or without (CTL) ASNase (0.003 IU/ml) for 24 hours. Data are expressed as mean ± SD (n= 4 experiments). **c.** Proliferation of WT and C1 Eµ-*Myc* cells treated as in **b.** until 72 hours. Data are expressed as the mean number of live cells (DAPI-) ± SD (n= 4 experiments). P-value from t-test. **d.** Total protein extracts prepared from whole-Eµ-*Myc* (#506) cells treated with ASNase as in **a.** at the indicated times and periods were immunoblotted for the indicated proteins. ERK2, loading control. Relative quantification below (normalized to the condition - ASNase of the 1^st^ challenge). Immunoblots shown are representative of 3 independent experiments. **e.** Schematic representation of the two successive therapeutic challenges with Vehicle (NaCl 0.9%) or ASNase in C57BL/6 mice bearing Eµ-*Myc* #506 lymphoma. The 1^st^ therapeutic challenge resulted in ^V^BCL and ^A^BCL of Vehicle- and ASNase-treated mice (n=6/group), respectively. Malignant B cells from axillary ^V^BCL or ^A^BCL, were then transferred into secondary recipient WT C57BL/6 mice. Seven days later, mice were treated with Vehicle or ASNase (2^nd^ therapeutic challenge). Resulting tumors are labelled Vehicle-^V^BCL, Vehicle-^A^BCL and ASNase-^V^BCL, ASNase-^A^BCL (n=6 mice/group). **f. g.** Survival curve of WT C57BL/6 mice intravenously injected with ^V^BCL (**f.**) or ^A^BCL (**g.**) malignant cells and treated with Vehicle or ASNase every 48 hours (n=6 mice/group). P value from log-rank test. **h.** Principal-component analysis (PCA) of metabolites abundance based on 130 metabolites detected by LC/MS in BCL of ^V^BCL cells or ^A^BCL cells-bearing C57BL/6 mice treated with Vehicle or ASNase (n=6 mice/group). **i.** Heatmap representation of asparagine and serine abundance in BCL presented in **e.** (n=6 mice/group). Malignant cells harvested from BCL of the Vehicle- and ASNase-treated Eµ-*Myc* cells-bearing C57BL/6 mouse #738 and #750, respectively, were engrafted into secondary recipient WT C57BL/6 mice for the 2^nd^ ASNase challenge. **j.** Total protein extracts prepared from BCL described in **i.** (^V^BCL, n=1; ^A^BCL, n=1; Vehicle ^A^BCL, n=6; ASNase-^V^BCL, n=6), were immunoblotted for the indicated proteins. ERK2, loading control. *ns*, not significant; **, p < 0.01; ***, p < 0.001; ****, p < 0.0001.

The expression levels of PHGDH, PSAT1, PSPH, ASNS and of the transcription factor ATF4, which were initially upregulated within the first 24 hours, remained elevated through 4 days of ASNase treatment (Figures 4d). Following therapy withdrawal, ASNS and PSPH protein levels returned to baseline within 24 hours, while PSAT1 and PHGDH levels modestly decreased at 72 hours and 96 hours (control condition of the 2^nd^ challenge), respectively, consistent with a time-dependent decreased expression level of ATF4. Upon ASNase re-exposure (2^nd^ challenge) these enzymes were re-induced within 24 hours (Figures 4d), confirming a dynamic and reversible regulatory process.

*In vivo*, Eµ-*Myc* cells isolated from BCL treated with either Vehicle or ASNase (1^st^ *in vivo* challenge) were re-engrafted into WT C57BL/6 mice and subjected to a second round of ASNase therapy (2^nd^ *in vivo* challenge). Mice engrafted with ASNase-treated cells still exhibited a significant survival benefit upon re-treatment (Figures 4e-f-g), supporting a model of flexible, non-permanent metabolic adaptation.

To assess whether the metabolic response to ASNase was reversible *in vivo*, we performed metabolomic profiling of BCL harvested after the 1^st^ or 2^nd^ ASNase treatment (*in vivo* challenge). Tumors from Vehicle-treated mice (Vehicle-^V^BCL and Vehicle-^A^BCL) exhibited distinct metabolic profiles compared to those from ASNase-treated mice (ASNase-^V^BCL and ASNase-^A^BCL), regardless of tumor origin (^V^BCL or ^A^BCL). Importantly, tumors harvested from Vehicle-treated mice (Vehicle-^V^BCL and Vehicle-^A^BCL) showed a highly similar metabolic signature, even though Vehicle-^A^BCL originate from ^A^BCL cells, i.e., malignant B cells that had already adapted to ASNase treatment, suggesting a transient resistance phenotype (Figure 4h). Among the most significantly deregulated metabolic pathways in ASNase-^A^BCL compared to Vehicle-^A^BCL, serine/glycine metabolism was prominent, consistent with previous findings (Figure S5c). ASNase-treated BCLs showed elevated serine and reduced asparagine levels, whereas the opposite pattern was observed in Vehicle-treated BCLs, underscoring dynamic metabolic reprogramming (Figure 4i). In line with our *in vitro* results (Figures 4d), induction of PSAT1, PSPH, and ASNS expression in BCL following 1^st^ *in vivo* challenge with ASNase, was transient, as their expression levels decreased in Vehicle-^A^BCL of Vehicle-treated mice (or was maintained in ASNase-^A^BCL of ASNase-treated mice) following the 2^nd^ *in vivo* challenge (Figures 4j and S5d). Collectively, our data highlight the metabolic plasticity of malignant B cells exposed to ASNase therapy.

### PHGDH activity limits ASNase-induced oxidative stress in malignant B cells

Given that amino acid or glucose starvation induces oxidative stress ^36, 37^, we determined whether ASNase treatment induces oxidative stress in Eµ-*Myc* cells and how these ASNase-sensitive Eµ-*Myc* cells adapt. Asparagine-deprived Eµ-*Myc* cells displayed elevated ROS levels and a higher oxidized to reduced glutathione ratio (GSSG/GSH) (Figures 5a-b and S6a). This oxidative stress was accompanied by NRF2 (Nuclear Factor Erythroid 2-Related Factor 2) protein stabilization that in turn activated the transcription of canonical antioxidant response genes, including Glutamate-cysteine ligase catalytic subunit (*Gclc*), Glutamate-cysteine ligase modifier subunit (*Gclm*) —the first rate-limiting enzymes of glutathione biosynthesis— as well as the NAD(P)H quinone dehydrogenase 1 (*Nqo1*) (Figures 5c-d-e).

**Figure 5:**
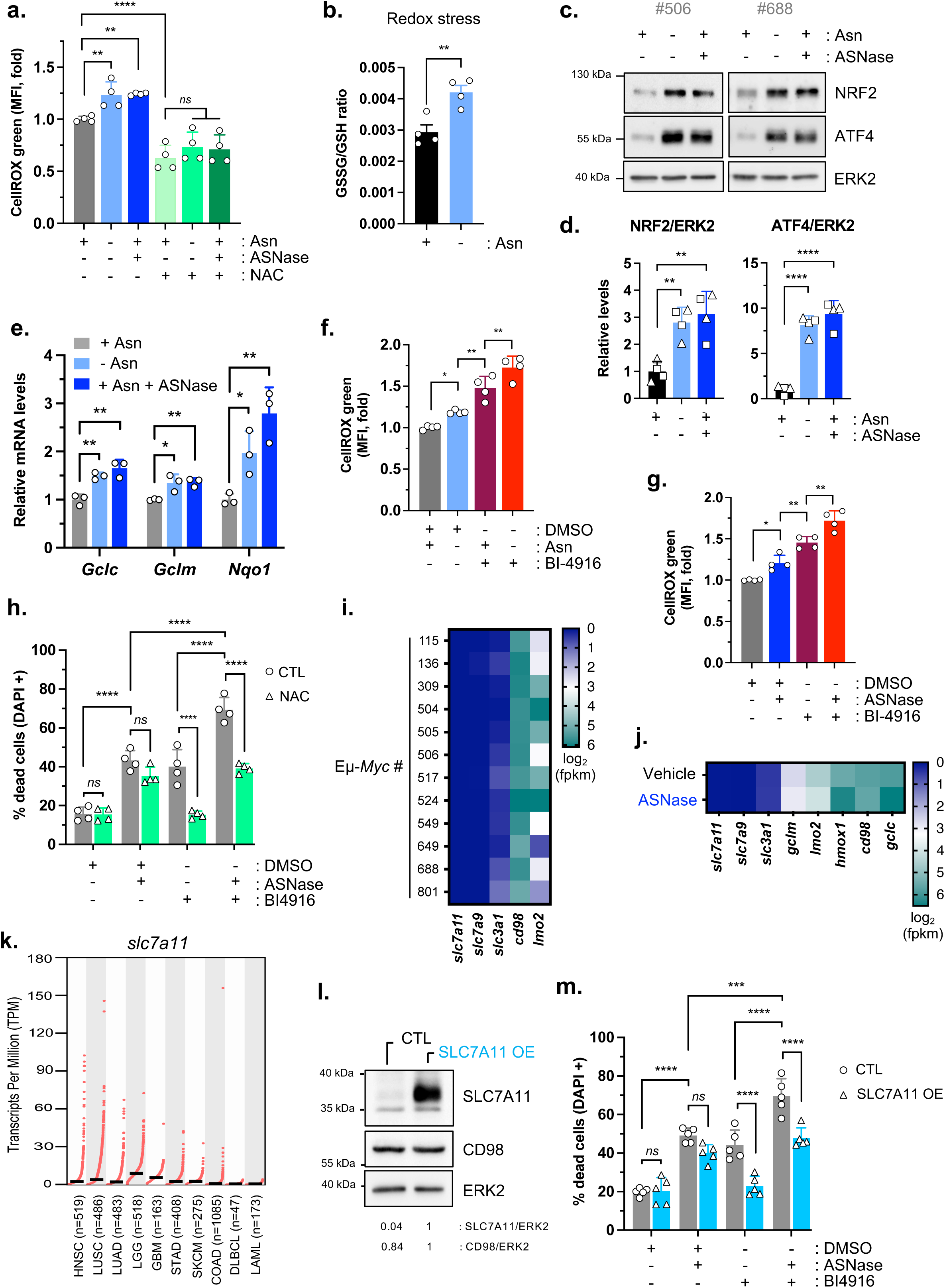
ASNase-treated malignant B cells activate an antioxidant defense program that depends on PHGDH activity to limit ROS production. **a.** Relative ROS levels in Eµ-*Myc* (#688) cells cultivated for 15 hours in Asn-free medium, supplemented with (+) or without (-) Asn (0.37 mM) and ASNase (0.003 IU/ml). Prior incubation with the CellROX probe, cells were pretreated (+) or not (-) with N-acetyl-L-cysteine (NAC, 10 mM) for 1 hour, followed by analysis by flow cytometry. Data are expressed as the Median Fluorescence Intensity (MFI) normalized to the control condition (+Asn) and presented as mean ± SD (n=4 experiments). P-value from 2-way Anova followed by Tukey’s test. **b.** GSSG/GSH ratio in Eµ-*Myc* (#688) cells incubated for 24 hours in Asn-free medium, supplemented (+) or not (-) with Asn (0.37 mM) for 24 hours. Data are expressed as mean ± SD (n= 4 biological replicates). P-value from t-test. **c.** Total protein extracts prepared from whole Eµ-*Myc* cells cultivated for 24 hours as described in **a.** were immunoblotted for NRF2 and ATF4. ERK2, loading control. Immunoblots shown are representative of 2 independent experiments for each Eµ-*Myc* line. **d.** Relative quantification of indicated protein levels expressed in Eµ-*Myc* #506 (square, n=2 independent experiments) and in Eµ-*Myc* #688 (triangle, n=2 independent experiments) cells. Data are normalized to the control condition +Asn and represented as mean ± SD. P value from t-test. **e.** Relative mRNA expression levels of *Gclc*, *Gclm* and *Nqo1* normalized to *Rplp0* in Eµ-*Myc* (#688) cells after 24 hours incubation under the conditions described in **a.** Data are expressed as mean ± SD (n= 3 experiments). P-value from t-test. **f. g.** Relative ROS levels in Eµ-*Myc* (#688) cells treated for 20 hours with (+) or without (-) Asn (0.37 mM) and/or the PHGDH inhibitor BI-4916 (10 µM) in Asn-free medium (**f.**) or with (+) or without ((-), DMSO) ASNase (0.003 IU/ml) and/or BI-4916 (10 µM) in Asn-containing medium (**g.**). Data are represented as MFI of CellROX green probe, normalized to control condition (+Asn) and expressed as mean ± SD (n= 4 experiments). P-value from 2-way Anova followed by Tukey’s test. **h.** Percentage of dead Eµ-*Myc* (#506) cells (DAPI+) treated with (+) or without (-) DMSO, ASNase (0.003 IU/ml), BI-4916 (7.5 µM) and/or NAC (5 mM) for 48 hours in Asn-containing medium. Data are expressed as mean ± SD (n= 4 experiments). P-value from 2-way Anova followed by Tukey’s test. **i.** Heatmap representation of mRNA expression levels for the indicated genes, determined by RNAseq from Eµ-*Myc* cells isolated from 12 individual transgenic Eµ-*Myc*^Tg/+^ mice. Expression values are shown as log_2_(fpkm). **j.** As in **i.** from BCL harvested from Eµ-*Myc* cells-bearing mice treated with Vehicle or ASNase (n=1 tumor/group). Expressions values are shown as log_2_(fpkm). **k.** *slc7a11* mRNA expression levels (transcript per million) in selected tumor entities available on the TCGA database. HNSC, Head and Neck squamous cell carcinoma; LUSC, Lung squamous cell carcinoma; LUAD, Lung adenocarcinoma; LGG, Brain Lower Grade Glioma; GBM, Glioblastoma multiforme; STAD, Stomach adenocarcinoma; SKCM, Skin Cutaneous Melanoma; COAD, colon adenocarcinoma; DLBCL, Diffuse Large B-cell Lymphoma; LAML, Acute Myeloid Leukemias. **l.** Whole-cell lysates prepared from Eµ-*Myc* (#506) cells stably expressing either control vector (CTL) or the murine SLC7A11-encoding vector (SLC7A11 OE), were immunoblotted for the indicated proteins. ERK2, loading control. Relative quantification below (normalized to the SLC7A11 OE condition). **m.** Percentage of dead control (CTL) and SLC7A11 overexpressing Eµ-*Myc* (#506) cells (DAPI+) treated with (+) or without (-) DMSO, ASNase (0.003 IU/ml) and/or BI-4916 (7.5 µM) for 48 hours in Asn-containing medium. Data are expressed as mean ± SD (n= 5 experiments). P-value from 2-way Anova followed by Tukey’s test. *ns*, not significant; *, p < 0.05; **, p < 0.01; ***, p < 0.001; ****, p < 0.0001.

Given the induction of *Gclc*, *Gclm* and the NRF2-targeted gene *slc6a9* (solute carrier family 6 member 9) ^38^, encoding a glycine transporter (Figure S6b), we verified whether glutathione synthesis, which requires cysteine, glutamate and ultimately glycine, was enhanced in asparagine-deprived Eµ-*Myc* cells. Using ^13^C_2_-Glycine, we detected increased levels of glycine-^13^C-labelled GSH (m+2) derived from ^13^C_2_-Glycine (Figures S6c-d), indicating active glutathione synthesis in asparagine-deprived Eµ-*Myc* cells. Of note, glycine appears to be specifically catabolized in glutathione biosynthesis, as evidenced by a significant reduction in the abundance of downstream products of glycine metabolism and diminished ^13^C-ATP (m+2) levels (Figure S6e).

Since *de novo* serine biosynthesis also contributes to glycine for antioxidant defense and given the observed increased levels of ^13^C_3_-serine and ^13^C_2_-glycine derived from [U-^13^C]-glucose in ASNase-treated cells (Figures 2f-g), we hypothesized that this pathway supports redox homeostasis. Inhibition of PHGDH with BI-4916 significantly increased ROS levels, indicating that *de novo* serine biosynthesis plays a critical role in maintaining redox homeostasis in Eµ-*Myc* cells under basal conditions (Figures 5f-g). Furthermore, cells treated with BI-4916/ASNase combination exhibit higher ROS levels compared to those observed with each single treatment, supporting the contribution of *de novo* serine biosynthesis to redox homeostasis in ASNase-treated Eµ-*Myc* cells. Accordingly, scavenging ROS with N-acetyl-L-cysteine (NAC) prevented cell death induced by BI-4916, either alone or in combination with ASNase (Figure 5h).

To promote an effective antioxidant response, cancer cells must not only meet a high demand for glycine but also cysteine, the rate-limiting precursor of glutathione biosynthesis ^39–41^. Due to the oxidizing tumor microenvironment, extracellular cysteine is unstable and rapidly converts into cystine. Therefore, most cancer cells rely on cystine transport, which is imported and reduced to cysteine in the cytosol, or on *de novo* cysteine biosynthesis. Surprisingly, Eµ-*Myc* cells harvested from 12 individual transgenic Eµ-*Myc* ^Tg/+^ mice, lacked detectable levels of s*lc7a11* mRNA, the solute carrier family 7 member 11 (also called xCT), which is the main plasma membrane antiporter mediating extracellular cystine uptake in exchange for glutamate (Figure 5i). *slc7a11* mRNA remained undetectable by qPCR in Eµ-*Myc* cells treated with ASNase for 72 hours *in vitro* or by RNAseq in BCL exposed to several weeks of ASNase treatment *in vivo* (Figures 5j and S6f). Other cystine transporters, *slc7a9* and *slc3a1,* were also absent, while *slc3a2* (also called CD98), encoding the chaperone for xCT and LATs transporters, was ubiquitously expressed (Figures 5i-j). Analysis of The Cancer Genome Atlas (TCGA) database confirmed that human DLBCL exhibit low *slc7a11* mRNA expression (Figure 5k), a finding reinforced by RNA-seq data from 775 human DLBCL samples ^42^, which showed that 7.28% expressed little to no detectable levels of *slc7a11 mRNA*, despite heterogeneous expression of *slc3a2* mRNA (Figures S6g-h). Overexpression of the murine SLC7A11 in *slc7a11*-deficient Eµ-*Myc* cells, attenuated the death induced by inhibition of PHGDH alone or in combination with ASNase (Figures 5l-m). Collectively, our results highlight the role of PHGDH in mitigating ASNase-induced oxidative stress in malignant B cells.

### ASNase-induced ROS revealed a therapeutic vulnerability to the clinically used PARP1/2 inhibitor in ASNase-sensitive malignant B cells *in vitro*

We next aimed to broaden our investigation into the possible consequences of perturbed redox homeostasis in ASNase-treated malignant B cells. Given that ROS accumulation can induce single-strand or double-strand DNA breaks, we sought to determine if ASNase-induced oxidative stress triggers DNA damage.

Asparagine-restricted Eµ-*Myc* cells exhibited increased phosphorylation of histone H2AX at Ser139 (γH2AX), along with activation of the ATR-dependent DNA damage response (DDR) pathway. This was evidenced by phosphorylation of ATR at Ser428 and its downstream substrates Chk1 at Ser345 and RPA32 at Ser33, both *in vitro* (Figures 6a-b-c-d) and *in vivo* (Figures S7a-b-c), indicating the presence of DNA damage. In contrast, the ATM-dependent DDR was not activated, as KAP1, a downstream substrate of ATM, was not phosphorylated at S^824^.

**Figure 6:**
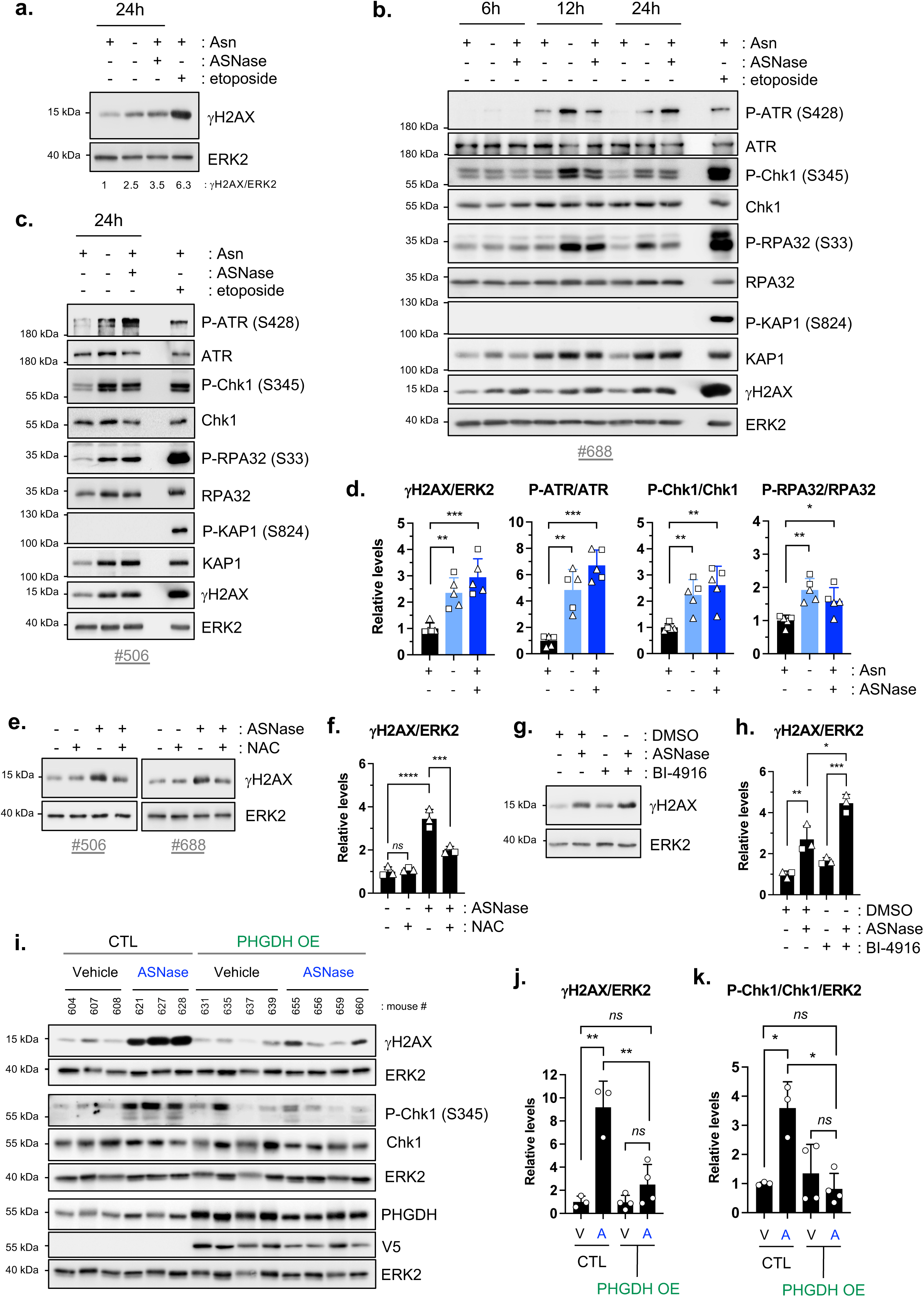
ASNase-treated malignant B cells exhibit oxidative stress-associated DNA damage, which are prevented by PHGDH overexpression. **a. b. c**. Total protein extracts prepared from whole Eµ-*Myc* #688 cells (**a.** and **b.**) or Eµ-*Myc* #506 cells (**c.**) cultured in Asn-free medium, supplemented (+) or not (-) with Asn (0.37 mM) and ASNase (0.003 IU/ml) for the indicated period, were immunoblotted for the indicated proteins. As a positive control, total protein lysates prepared from whole Eµ-*Myc* cells treated with etoposide (1µg/ml) for 3 hours were used. ERK2, loading control. (**a.**) Relative quantification below (normalized to the control condition +Asn). **d.** Relative quantification of the expression levels of indicated proteins in Eµ-*Myc* #688 cells (triangle, n=3 independent experiments) and in Eµ-*Myc* #506 cells (square, n=2 independent experiments), following 24 hours of incubation as in **a.** Data are represented as mean ± SD. P-value from 2way Anova, followed by Tukey’s test. **e.** γH2AX expression in Eµ-*Myc* cells (#506 and #688) following 24 hours incubation in Asn-containing medium supplemented (+) or not (-) with N-acetyl-L-cysteine (NAC, 10 mM) and/or ASNase (0.003 IU/ml). ERK2, loading control. Immunoblots shown are representative of 3 independent experiments. **f.** Relative quantification of γH2AX expression in Eµ-*Myc* cells presented in **e.** (#688, triangle, n=2 independent experiments; #506, square, n=1 experiment). Data are represented as mean ± SD. P-value from one-way Anova, followed by Tukey’s test. **g.** γH2AX expression in Eµ-*Myc* (#688) cells treated (+) or not (-) for 24 hours with DMSO (-/-), ASNase (0.003 IU/ml) and/or BI-4916 (10 µM). ERK2, loading control. Immunoblots shown are representative of 3 independent experiments. **h.** Relative quantification of γH2AX expression in Eµ-*Myc* cells presented in **g.** (#688, triangle, n=2 independent experiments; #506, square, n=1 experiment). Data are represented as mean ± SD. P-value from 2way Anova, followed by Tukey’s test. **i.** Total protein extracts prepared from B-cell lymphomas harvested from C57BL/6 mice engrafted with control (CTL) or V5-tagged murine PHGDH-overexpressing (PHGDH OE) Eµ-*Myc* (#506) cells and treated with Vehicle or ASNase for 5 weeks until disease endpoint, were immunoblotted for the indicated proteins (CTL-Vehicle and CTL-ASNase, n=3 mice/group; PHGDH OE-Vehicle and PHGDH OE-ASNase, n=4 mice/group). ERK2, loading control. **j. k.** Relative quantification of γH2AX (**j.**) and P-Chk1 (Ser345) (**k.**) expressed in BCL presented in **i.** Data are normalized to the control condition CTL-Vehicle and represented as mean ± SD. P-value from 2way Anova, followed by Tukey’s test. V: Vehicle; A: ASNase. *ns*, not significant; *, p < 0.05; **, p < 0.01; ***, p < 0.001; ****, p < 0.0001.

Scavenging ROS with NAC reduced γH2AX levels, confirming that oxidative stress contributes to ASNase-induced DNA damage (Figures 6e–f). Conversely, PHGDH inhibition modestly increased γH2AX in ASNase-treated cells (Figures 6g–h). Consistently, PHGDH overexpression in Eµ-*Myc* cells (Figures S4l-m) significantly attenuated γH2AX expression levels and activation of the ATR-dependent DDR during ASNase therapy *in vivo* (Figures 6i-j-k). This supports a protective role for *de novo* serine biosynthesis in preserving genome stability.

Following an ATR-dependent DNA damage response, the chromatin-associating enzymes poly (ADP-ribose) polymerase 1 (PARP1) and PARP2 catalyze the synthesis and transfer of ADP-ribose polymer (PARs) from nicotinamide adenine dinucleotide (NAD^+^) onto their serine, tyrosine, and glutamate residues (auto-PARylation), activating them. In turn, activated PARPs enables the PARylation of target proteins, promoting their recruitment to DNA breaks for effective DNA repair. We aimed to investigate whether PARP was activated in ASNase-treated Eµ-*Myc* cells, as this could represent a therapeutically exploitable tumor vulnerability using clinically approved PARP1/2 (PARP) inhibitors. We observed increased expression of proteins harboring poly(ADP-ribose) motifs, indicating elevated PARP activity in asparagine-restricted Eµ-*Myc* cells (Figure 7a). PARP inhibitors, such as Olaparib, compete with NAD^+^ for the catalytically active site of PARP, thereby preventing the initiation of DNA repair mechanisms. To interfere with DNA repair in ASNase-treated cells, we sought to test the effects of ASNase/Olaparib co-treatment in Eµ-*Myc* cells, in *in vitro* and *in vivo* settings. This therapeutic combination enhanced caspase-dependent cell death in two OxPhos-dependent Eµ-*Myc* lines (#506 and #688), surpassing the effect of either agent alone (Figures 7b-c-d). When co-treated with Olaparib (0.3 µM), ASNase-treated Eµ-*Myc* cells did not resume proliferation *in vitro* (Figure 7e-f), even after 14 days (data not shown). *In vivo*, co-treatment with ASNase (every 48 hours) and daily Olaparib significantly delayed BCL onset and weight loss in C57BL/6 mice, compared to monotherapies (Figures 7g-h-i). ASNase/Olaparib co-treatment antineoplastic effect was confirmed in a human colon cancer cell line grown in 3D (Figures S8a-b). Overall, these results demonstrate that ASNase-induced oxidative stress creates a vulnerability to the PARP inhibitor Olaparib in malignant B cells, and that combining ASNase with Olaparib holds potential as an effective anti-cancer strategy.

**Figure 7:**
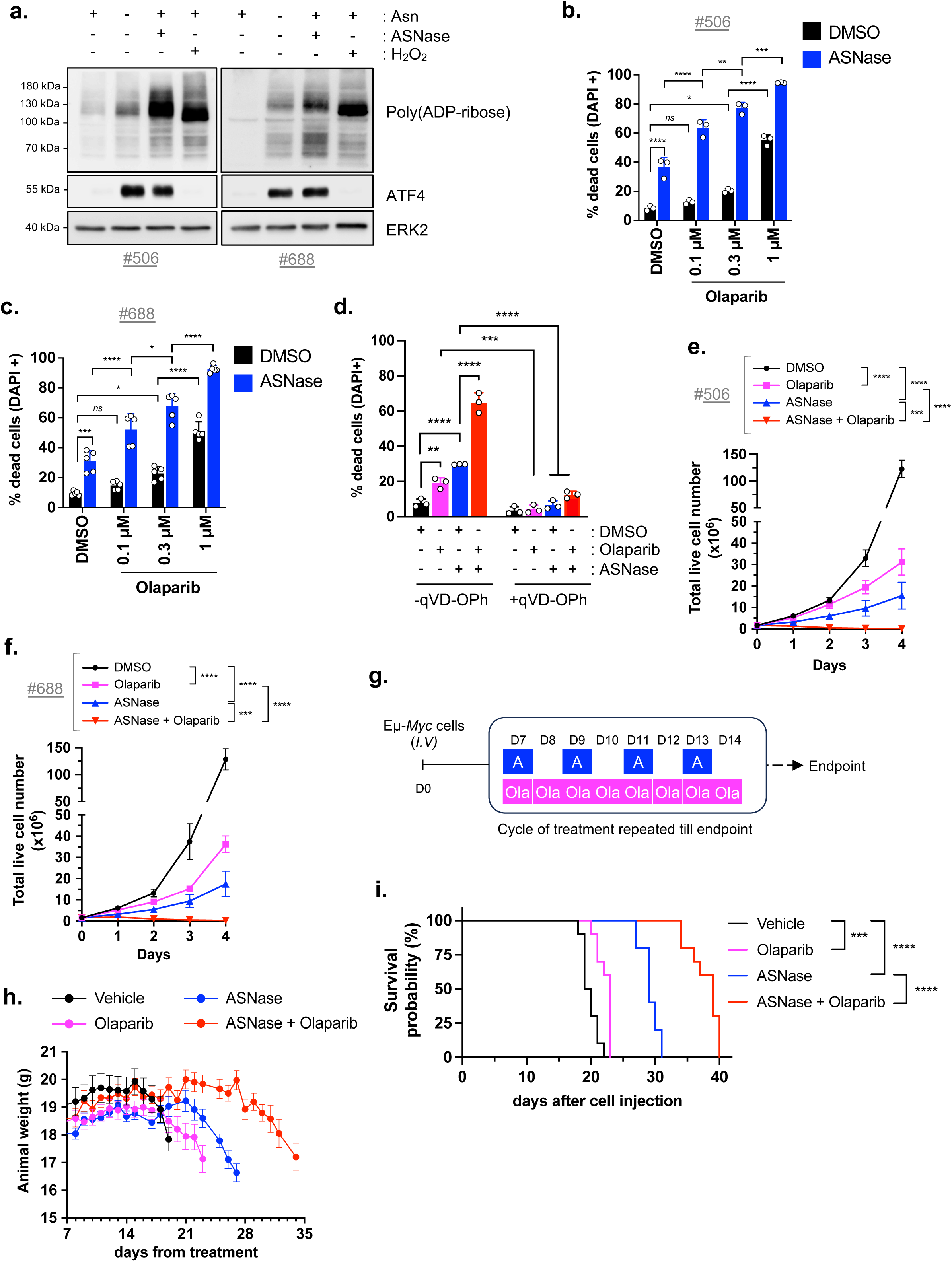
DNA damage in ASNase-treated malignant B cells revealed their sensitivity to the PARP1/2 inhibitor, Olaparib. **a.** Total protein extracts prepared from whole Eµ-*Myc* cells (#506 or #688) cultivated for 10 hours in Asn-free medium, supplemented (+) or not (-) with Asn (0.37 mM) alone or in combination with ASNase (0.003 IU/ml), were immunoblotted using an anti-poly(ADP-ribose) antibody. As a positive control, total protein lysates prepared from whole Eµ-*Myc* cells stimulated with H_2_O_2_ (0.5 mM) for 5 minutes were used. ERK2, loading control. Immunoblots shown are representative of 2 independent experiments for each Eµ-*Myc* line. **b. c.** Percentage of dead Eµ-*Myc* (#506) cells (**b.**) or Eµ-*Myc* #688 cells (**c.**) (DAPI+) treated with DMSO, *E-Coli* L-asparaginase (ASNase, 0.003 IU/ml) and/or the PARP inhibitor Olaparib (at indicated concentrations) for 24 hours in Asn-containing medium. Data are expressed as mean ± SD (**b.**, n=3 experiments; **c.**, n=5 experiments). P-value from 2-way Anova, followed by Sidak’s test. **d.** Percentage of dead Eµ-*Myc* (#688) cells (DAPI+) treated with DMSO, ASNase (0.003 IU/ml) and/or Olaparib (0.3 µM) for 24 hours in Asn-containing medium and in the presence (+) or absence (-) of the caspase inhibitor qVD-OPh (20 µM). Data are expressed as mean ± SD (n=3 experiments). P-value from 2way Anova, followed by Tukey’s test. **e. f.** Four-days proliferation of Eµ-*Myc* (#506) cells (**e.**) and of Eµ-*Myc* (#688) cells (**f.**) treated as described in **d.** Data are expressed as the mean number of live cells (DAPI-) ± SD (n=4 experiments). P-value from 2way Anova, followed by Tukey’s test. **g.** Schematic representation of *in vivo* co-treatment with ASNase/Olaparib. **h.** Body weight of Eµ-*Myc* (#506) cells-bearing C57BL/6 mice from the start of treatment (day 7 post tumor cell inoculation) (n=10 mice/group). **i.** Survival curves of WT C57BL/6 mice intravenously injected with Eµ-*Myc* (#506) cells and treated 7 days later with Vehicle, Olaparib (daily), ASNase (every 48 hours) or their combination, for several weeks, until lymphoma development reached endpoint (n=10 mice/group). P-value from log-rank test. *ns*, not significant; *, p < 0.05; **, p < 0.01; ***, p < 0.001; ****, p < 0.0001.

## Discussion

Our study reveals that the clinically used ASNase treatment induces oxidative stress and associated DNA damage in malignant B cells, therefore triggering a compensatory reliance on PHGDH-driven serine biosynthesis to restore redox stress and on PARP-mediated DNA repair to resolve the damage. Pharmacological inhibition of PARP with Olaparib significantly sensitizes malignant B cells to ASNase, confirming the functional importance of this adaptive response. This metabolic-DNA damage crosstalk not only supports tumor cell survival under treatment-induced nutritional stress but also exposes a therapeutically actionable liability. Our findings thus provide a strong rationale for combinatorial strategies targeting PARP in ASNase-treated malignant B cells using clinically approaved PARP inhibitors, to improve the efficacy and durability of ASNase-based therapies.

Most studies investigating mechanisms of resistance to ASNase treatment in solid and hematological cancer cell lines have focused on primary resistance, aiming to identify novel therapeutic strategies to eliminate ASNase-resistant cancer cells ^10, 11, 18–20, 43–46^. Several CRISPR-Cas9 screens have identified determinants of ASNase sensitivity in long-term ASNase-exposed resistant cell lines (10–20 days *in vitro*) ^43, 45, 46^ or *in vivo* in immunodeficient mice ^20^. One such screen highlighted the overexpression of PHGDH, PSAT1 and PSPH, the three enzymes involved in *de novo* serine biosynthesis, in ASNase-resistant melanoma cells knocked-out for the asparagine synthetase ^45^. However, whether *de novo* serine biosynthesis is functionally enhanced in resistant cells and whether its enzymes drive resistance remained not established. Using ASNase-sensitive B-cell lymphomas, our study identified novel determinants of tumor relapse during ASNase therapy, following an initial antineoplastic response, resembling secondary resistance as observed in R/R relapsed DLBCL patients ^8^. We observed elevated levels of *de novo*-synthesized serine and increased oxidative stress in ASNase-treated malignant B cells. We identified PHGDH as a key mediator of this adaptive response by mitigating ROS accumulation, thereby enabling the outgrowth of therapy-stressed malignant B cells. These findings are consistent with a recent study showing that Eµ-*Myc*-driven lymphomas rely on *de novo* serine biosynthesis for disease progression, highlighting PHGDH as a clinically relevant target in human B-cell lymphomas ^47^. Given that the asparagine synthetase is a key determinant in ASNase resistance, it remains to be addressed whether activation of *de novo* serine biosynthesis in ASNase-treated cells depends on their ability to replenish sufficient asparagine levels through upregulation and/or activation of asparagine synthetase on treatment *in vitro* and *in vivo*.

Interestingly, cycles of ASNase therapy and drug withdrawal in chemo-naïve malignant B cells, demonstrated that their progeny was still sensitive to rechallenge with therapy, a transient tumor adaptation observed *in vitro* and *in vivo*. Moreover, this tumor-intrinsic adaptative response involved a flexible metabolic remodeling, which engage *de novo* serine, glycine, glutathione, and asparagine biosynthesis, likely driven by a dynamic ATF4 and NRF2-dependent transcriptional program that counteracts ASNase-induced oxidative stress and facilitates cell cycle re-entry. Consistent with our results, asparagine restriction induces oxidative stress and an NRF2-dependent stress response in CD_8_^+^ T cells, enhancing their proliferation and effector functions ^48–51^.

We also demonstrated that asparagine deprivation induces oxidative stress and DNA damage. ASNase-treated malignant B cells activated the ATR-dependent DNA damage response pathway and exhibited elevated PARP activity, suggesting replication stress. Consequently, ASNase-treated BCL are highly sensitive to the PARP inhibitor Olaparib, both *in vitro* and *in vivo*, implicating PARP as a determinant of BCL relapse during ASNase treatment. Our findings are consistent with prior studies showing that PARP plays a key role in DNA repair mechanisms following hydrogen peroxide-induced single-strand DNA breaks ^52^ and for maintaining genomic integrity in B cells ^53^. Notably, PARP2 loss increases replication stress and impedes BCL progression in the Eµ-*Myc* model ^53^.

Overall, our study revealed a crosstalk between amino acids availability, redox metabolism, and genome integrity, with PHGDH and PARP as critical regulators. In line with our findings, the NRF2-targeted glucose-6-phosphate dehydrogenase (G6PD), was recently shown to regulate NAD metabolism and basal PARylation, linking redox control to DNA replication integrity in p53-mutant colorectal cancer cells ^54^. Whether ASNase-induced metabolic reprogramming influences NAD⁺ biosynthesis to regulate PARP activation and DNA repair mechanisms in malignant cells warrants further investigations.

Importantly, human ALL samples resistant to ASNase treatment displayed enrichment of genes involved in double-strand break repair via break-induced replication 55, reinforcing our observation that asparagine preserves genome stability in malignant cells.

A key feature of PARP inhibitors is their effectiveness in targeting solid cancers with defects in homologous recombination (HR), most often due to germline or somatic BRCA1/2 mutations. HR deficiency is prevalent in advanced breast, ovarian and metastatic pancreatic cancers, and is a key factor predicting tumor response to PARP inhibition ^56, 57^. Although BRCA1/2 mutations are rare in hematological malignancies, a recent study provided strong rationale for PARP inhibitor use in DLBCL subsets with HR deficiency. Specifically, PARP inhibition was shown to synergize with anti-CD20-based chemotherapies in a subtype of human DLBCL characterized by LIM-domain 2 (LMO2)-driven homologous recombination deficiency, a feature not observed in DLBCL lacking LMO2 ^58^. LMO2 is a cysteine-rich protein preventing BRCA1 recruitment to DNA-strand breaks ^58^. Malignant Eµ-*Myc* cells used in our study express variable levels of *lmo2* mRNA (figures 5i-j), which may, in part, explain their sensitivity to Olaparib treatment, particularly in combination with ASNase. Further investigations would be needed to comprehensively characterize the HR status of Eµ-*Myc* lymphomas and the mechanisms regulating it. Nevertheless, the antineoplastic effect of ASNase/Olaparib co-treatment was confirmed in HR-proficient human colorectal cancer cells (HCT-116) grown in 3D, suggesting that other HR-proficient cancers may benefit from this combination strategy. Assessing toxicity in non-cancerous cells (e.g., immune cells, hepatocytes, cardiomyocytes) or normal tissues in treated WT mice would require dedicated pharmacological studies to ensure cancer cell-specific effects. That said, Olaparib is well-tolerated in normal cells with functional HR ^59^, possibly offering a therapeutic window in which malignant cells would be preferentially targeted by ASNase/Olaparib co-treatment.

Overall, our study sheds light on the dynamic and flexible tumor metabolic response during ASNase therapy. These insights not only enhance our understanding of the metabolic mechanisms facilitating resistance to a clinically approved drug, but also unveil clinically actionable combination strategy to enhance the therapeutic index of ASNase and tackle aggressive B-cell lymphomas by combining ASNase and Olaparib. While PARP inhibitors are currently being evaluated in combination with radiation, alkylating agents, topoisomerase I inhibitors, PI3K inhibitors, and immunotherapies ^60^, our findings establish a proof-of-concept that PARP inhibition can be effectively combined with a metabolic drug—ASNase—in B-cell lymphomas, particularly those expressing LMO2. Collectively, our findings revealed a clear path for developing an anti-tumor strategy that combines ASNase and PARP1/2 inhibitor, offering a new therapeutic opportunity for patients treated with ASNase or Olaparib, to prevent tumor relapse.

## Materials and Methods

### Mice

C57BL/6 Eµ-*Myc* transgenic mice (Eµ-*Myc*^Tg/+^) were purchased from the Jackson Laboratory (002728) and are bred and maintained at our local animal facility (C3M, INSERM U 1065, Nice, France, B 06-088-20). Wild-type (WT) C57BL/6JOlaHsd female were purchased from Envigo and housed at our local animal facility. All mice were maintained in specific pathogen-free conditions and all experimental procedures were performed in compliance with the protocols approved (n° APAFIS#25228-2020032917261458 v9) by the Institutional Animal Care and Use Committee (IACUC of University Nice, Côte d’Azur) and the Ministère de la Recherche et de l’Innovation.

### Cell lines

Primary OxPhos-dependent Eμ-*Myc* #506 and Eμ-*Myc* #688 cells were isolated from different Eμ-*Myc*^Tg/+^ mice (mouse #506 and #688, respectively), as previously described and characterized ^8, 61^. These cells were maintained in DMEM-GlutaMAX medium (31966047, Thermo Fisher Scientific) supplemented with 10% of fetal bovine serum (FBS) (F7524, Sigma), 50 µM of 2-Mercaptoethanol (31350010, Thermo Fisher Scientific), 0.37 mM of L-asparagine (A0884, Sigma), 10 mM of HEPES pH 7.4 (15630056, Thermo Fisher Scientific) and 100 U/ml of Penicillin-Streptomycin (15140122, Thermo Fisher Scientific). 293T cells (CRL-1573, ATCC) were maintained in DMEM-GlutaMAX media (31966047, Thermo Fisher Scientific) supplemented with 10% FBS and 100 U/ml of Penicillin-Streptomycin and used to produce VSV-G retroviral vectors. The murine Lewis Lung carcinoma (LLC) cell line (CRL1642, ATCC) was maintained in DMEM-GlutaMAX medium (31966047, Thermo Fisher Scientific) supplemented with 10% FBS and 100 U/ml of Penicillin-Streptomycin. The murine colon carcinoma cell line CT26 (CRL-2638, ATCC) was maintained in RPMI-GlutaMAX medium (61870044, Thermo Fisher Scientific) supplemented with 10% FBS, 100 U/ml of Penicillin-Streptomycin and 1 mM of sodium pyruvate (11360070, Thermo Fisher Scientific). The murine colon adenocarcinoma cell line MC38 (SCC172, Sigma) was maintained in DMEM-F12-GlutamaX medium (31331028, Thermo Fisher Scientific) supplemented with 10% FBS, 1 mM of sodium pyruvate and 100 U/mL of Penicillin-Streptomycin. The human colon carcinoma cell line HCT116 (CVCL_0291) was obtained from ATCC, routinely cultured in McCoy medium (16600082, Thermo Fisher Scientific), supplemented with 10% FBS and 100 U/ml of penicillin/streptomycin. All cell lines are mycoplasma-free and routinely tested for mycoplasma.

### Media for *in vitro* studies

Asparagine-free medium refers to DMEM high D-glucose (4.5 g/L), no L-glutamine, no pyruvate medium (11960044, Thermo Fisher Scientific), supplemented with 10% FBS (F7524, Sigma), 3 to 50 µM of 2-Mercaptoethanol, 10 mM of HEPES pH 7.4, 100 U/ml of Penicillin-Streptomycin, 1 mM of sodium pyruvate (11360070, Thermo Fisher Scientific), 2 mM of L-glutamine (25030081, Thermo Fisher Scientific). This medium contains supraphysiological concentration of serine and glycine.

Asparagine-containing medium refers to the asparagine-free medium described above, supplemented with L-asparagine (0.37 mM).

Asparagine, serine, glycine-free medium refers to MEM no L-glutamine medium (21090022, Thermo Fisher Scientific), supplemented with 10% FBS (F7524, Sigma) dialyzed (132724, Spectra/Por®3 Dialysis Membrane Standard RC Tubing, MWCO 3.5 kDa), 50 µM of 2-Mercaptoethanol, 10 mM of HEPES pH 7.4, 100 U/ml of Penicillin-Streptomycin, 20 mM of D-glucose (J60067, Thermo Fisher Scientific), 1 mM of sodium pyruvate, 2 mM of L-glutamine, 1% of vitamins cocktail (11120052, Gibco).

Glucose and asparagine-free medium refers to DMEM no L-glutamine, no pyruvate medium (A14430-01, Thermo Fisher Scientific), supplemented with 10% FBS (F7524, Sigma), 50 µM of 2-Mercaptoethanol, 10 mM of HEPES pH 7.4, 100 U/ml of Penicillin-Streptomycin, 1 mM of sodium pyruvate, 2 mM of L-glutamine. This medium contains supraphysiological concentration of serine and glycine.

### ASNase formulations

For all *in vitro* experiments, the native *Escherichia-coli* L-asparaginase II (ASNase) (Kidrolase®, Jazz Pharmaceutical) was used. For *in vivo* studies presented in figures 1, 2, 4, S3 and S5, we used Kidrolase® and for *in vivo* studies in figures 3, 6, 7 and S3, we used the native *Escherichia-coli* L-asparaginase II recombinant protein (Spectrila®).

### *In vivo* transfer of Eµ*-Myc* cells and survival experiments

The transfer of mouse primary Eµ-*Myc* cells was performed into syngeneic, nontransgenic, 6-weeks-old C57BL/6JOlaHsd females (Envigo) by intravenous (*I.V*) injection of 100.000 live cells per recipient mouse. Seven days later, mice were treated with either Vehicle (NaCl 0.9%) or 2500 IU/kg of the native (Kidrolase®, Jazz Pharmaceutical) or recombinant (Spectrila®) *E-coli* L-asparaginase (ASNase), every 48 hours, by intraperitoneal (*I.P*) injection, until lymphoma reached the ethical limit. ASNase is diluted in NaCl 0.9%. Tumor onset was determined by inguinal lymph node palpation. Food was given *ad libitum*. No randomization was performed. At endpoint, ASNase-treated mice received a last ASNase injection and were sacrificed three to four hours after. When disease endpoint was reached, all mice bearing Eµ-*Myc* B-cell lymphomas (BCL) were sacrificed by cervical dislocation. Blood was then collected through cardiac punction, placed in lithium heparinate coated tubes (BD Microtainer), centrifuged 6000g for 2 min to isolate the plasma. The plasma samples were collected, snap-frozen and stored at -80°C for metabolomic analysis. To study the survival of mice engrafted with Eµ-*Myc* cells, previous experiments have shown statistical significance starting from 6 mice/group ^9^, a minimal number required to detect significant biological effects between two (or more) experimental groups, as confirmed using the G*power software. This number was confirmed using G power software Mice survival defined as survival probability (%) was determined as the time between the *I.V* injection of Eµ-*Myc* cells and the time when mice had to be sacrificed for ethical reasons when lymphomas progression reached the endpoint. Tumor-free mice (%) was determined as the time between the *I.V* injection of Eµ-*Myc* cells and the time when one inguinal lymphoma was palpable.

For the second *in vivo* therapeutic challenge, syngeneic, nontransgenic, 6-weeks-old C57BL/6JOlaHsd females (Envigo) were intravenously (*I.V*) injected with 100.000 live Eµ-*Myc* cells per recipient mouse. Eµ-*Myc* cells used were harvested from BCL obtained after the 1^st^ in vivo therapeutic challenge. Seven days later, mice were treated with Vehicle or ASNase at same concentration, frequency, and administration route as mentioned above.

For *in vivo* co-treatment with ASNase and Olaparib, syngeneic, nontransgenic, 6-weeks-old C57BL/6JOlaHsd females (Envigo) were intravenously injected with 100.000 live Eµ-*Myc* cells per recipient mouse. Seven days later, mice were treated with either Vehicle or 50 mg/kg of Olaparib (daily *I.P*) or 2500 IU/kg of Spectrila (*I.P* every 48 hours) or the combination of both drugs (daily *I.P* Olaparib 50 mg/kg and Spectrila, 2500 IU/kg, *I.P* every 48 hours). Olaparib (HY-10162, MCE) was dissolved in DMSO at a concentration of 100 mg/ml and then diluted at 5 mg/ml in 20% sulfobutylether-β-cyclodextrin solution (HY-17031, MCE, diluted in NaCl 0.9%). Vehicle consists in daily *I.P* injections of a 5% DMSO/95% sulfobutylether-β-cyclodextrin (in 20% in NaCl 0.9%) solution and/or *I.P* of NaCl 0.9% every 48 hours.

Food was given *ad libitum*. No randomization was performed. Endpoint was evaluated by blinding. Survival functions were estimated using the Kaplan-Meier method and compared with the log-rank test.

### RNA extraction, and Real-time Quantitative PCR

Total RNA was extracted from Eµ-*Myc* cells using the RNeasy mini kit (74104, Qiagen) according to manufacturer’s instructions. Reverse transcription was performed from 2 µg of RNA using the Omniscript RT kit (205113, Qiagen). Relative mRNA levels were obtained by real-time quantitative PCR (qPCR) on the StepOne™ Real-Time PCR System (Thermo Fisher Scientific) using the TaqMan assay primer set (Thermo Fisher Scientific) for murine *Phgdh* (Mm01623589_g1), *Psat1* (Mm01613328_g1), *Psph* (Mm01197775_m1), *Gls* (Mm01257297_m1), *Gclc* (Mm00802658_m1), *Gclm* (Mm01324400_m1), *Nqo1*(Mm01253561_m1), *slc7a11* (Mm00442530_m1), *slc6a9* (Mm00433662_m1), *Asns* (Mm01137310_g1), and the TaqMan Universal PCR Master Mix (4304437, Fisher Scientific) according to manufacturer’s instructions. The qPCR was performed from 8 ng of cDNA, except for *slc7a11* amplification for which we used 100 ng of cDNA. Relative mRNA levels were normalized to mouse *Rplp0* (Mm00725448_s1, Thermo Fisher Scientific) and to the control condition (+ Asn, 0.37 mM).

### RNAseq data

RNA sequencing (RNA-seq) alignment files and clinical information from the EGAS00001002606 study ^42^ were downloaded from the European Genome-phenome Archive. Quantification of genes was performed using featureCounts v1.5.0-p3 based on Ensembl GTF release 75 annotations. RNA-seq data were then normalized using the Bioconductor package DESeq2. Eighteen (out of 775) samples showing no expression of *Rplp0* were removed from the analysis.

RNA-seq data on human samples (solid cancers and hematological cancers) were retrieved and downloaded from The Cancer Genome Atlas program (TCGA, https://portal.gdc.cancer.gov/) through the Gene Expression Profiling Interactive Analysis website (http://gepia.cancer-pku.cn). The levels of *slc7a11* mRNA were represented in transcript per million (TPM).

The RNA sequencing from Eµ-*Myc* cells (*in vitro* and *in vivo*) was performed on Illumina Hiseq2000 at the IPMC functional genomics platform (France). Libraries were generated using Truseq Stranded mRNA kit (Illumina), from 1µg of total RNA extracted from different Eµ-*Myc* cell lines isolated from 12 individual transgenic Eµ-*Myc*^Tg/+^ mice and from B-cell lymphomas harvested from mice engrafted with Eµ-*Myc* #506 cells, and treated Vehicle or ASNase for several weeks. Libraries were then quantified with Qubit dsDNA High Sensitivity Assay Kit (Invitrogen) and pooled. 4nM of this pool were loaded on a Nextseq 500 High output Flowcell and sequenced on a NextSeq 500 platform (Illumina) with 2 × 75bp paired-end chemistry. Analysis of the obtained libraries sequences (reads) were performed with DESeq2-Bioconductor after assessing quality of the data and gene counts and normalization.

### DNA constructs

To amplify mouse *Asns* or *Phgdh* cDNA, total RNA was extracted from Eµ-*Myc* cells and reverse transcription was performed as mentioned above. The murine *Asns* coding sequence tagged to V5 was amplified by PCR from the resulting cDNA and cloned into the retroviral GFP-encoded pMIG vector (9044, Addgene) to produce the pMIG-*Asns*-v5 vector, as previously described ^9^. The murine *Phgdh* coding sequence (without stop codon) was amplified by PCR using a recombinant *Taq* DNA polymerase (10342020, Thermo Fisher Scientific) according to manufacturer’s instructions and with the following primers: forward 5’-GCAATGGCCTTCGCAAATCTGC-3’ and reverse 5’-GGAGCCAGCCCAGGCGT-3’. The PCR product was cloned into the pCR™ 2.1-TOPO® TA vector using the TOPO™ TA Cloning™ Kit (K455040, Thermo Fisher Scientific) according to manufacturer’s instructions. The resulting plasmid TOPO-*Phgdh* was amplified in One Shot TOP10 chemically competent *E.Coli* (C404003, Thermo Fisher Scientific) with ampicillin (A9518, Sigma) selection, purified (740414.50, Macherey-Nagel) and sequenced to confirm the cloning of the murine *wild-type Phgdh* coding sequence. Once confirmed, *Phgdh* coding sequence tagged to V5 was subcloned from the TOPO-*Phgdh* vector into the retroviral GFP-encoded pMIG vector (9044, Addgene), by PCR with the following primers: forward 5’-GAGAGTCGACATGGCCTTCGCAAATCTGC-3’ and reverse 5’-GGTTAACTCACGTAGAATCGAGACCGAGGAGAGGGTTAGGGATAGGCTTACCGAAGCAGAACTGGAAAGCCTCCAATAC-3’. Then, the amplicon was digested using SalI (R3136S, New England Biolabs) and HpaI (R0105S, New England Biolabs) restriction enzymes. The pMIG vector was linearized using XhoI (R0146S, New England Biolabs) and HpaI restrictions enzymes to generate compatible cohesive ends. Digested pMIG vector was dephosphorylated with Antarctic phosphatase (MO289S, Biolabs). The following double-strand DNA fragment which contains the coding sequence of the murine *slc7a11* with BglII and EcoRI restriction site was synthesized (Thermo Fisher Scientific): “ GGGGAGATCTATGGTCAGAAAGCCAGTTGTGGCCACCATCTCCAAAGGAGGTTACCTGCAGGGCAATATGAGCGGGAGGCTGCCCTCCATGGGGGACCAAGAGCCACCTGGGCAGGAGAAGGTAGTTCTGAAAAAGAAGATCACTTTGCTGAGGGGGGTCTCCATCATCATCGGCACCGTCATCGGATCAGGCATCTTCATCTCCCCCAAGGGCATACTCCAGAACACGGGCAGCGTGGGCATGTCCCTGGTTTTCTGGTCTGCCTGTGGAGTACTGTCACTTTTTGGAGCCCTGTCCTATGCAGAATTAGGTACAAGCATAAAGAAATCTGGTGGTCATTACACATACATTCTGGAGGTCTTTGGTCCTTTGCTGGCTTTTGTTCGAGTCTGGGTGGAACTGCTCGTAATACGCCCTGGAGCTACTGCTGTGATATCCCTGGCATTTGGACGCTACATCCTGGAACCATTTTTTATTCAATGTGAAATTCCTGAACTTGCAATCAAGCTCGTGACAGCTGTGGGCATCACTGTGGTGATGGTCCTAAATAGCACGAGTGTCAGCTGGAGTGCCCGGATCCAGATTTTCCTAACCTTTTGCAAGCTCACAGCAATTCTGATAATTATAGTCCCTGGAGTTATACAGCTAATTAAAGGGCAAACACATCACTTTAAAGATGCATTTTCAGGAAGAGACACAAGTCTAATGGGGTTGCCCTTGGCTTTTTATTATGGGATGTATGCATATGCTGGCTGGTTTTACCTCAACTTTATTACTGAAGAAGTAGACAACCCTGAAAAAACCATCCCCCTTGCAATCTGCATCTCCATGGCTATCATCACAGTGGGCTACGTACTGACAAACGTGGCCTATTTTACCACCATCAGTGCGGAGGAGCTGCTGCAGTCCAGCGCCGTGGCGGTGACCTTCTCTGAGCGGCTGCTGGGAAAATTCTCATTAGCAGTCCCGATCTTTGTTGCCCTCTCCTGCTTCGGCTCCATGAACGGTGGTGTGTTCGCTGTCTCCAGGTTATTCTACGTCGCATCTCGAGAAGGGCACCTTCCGGAAATCCTCTCTATGATTCATGTCCACAAGCACACTCCTCTGCCAGCTGTTATTGTTTTGCATCCTCTGACGATGGTGATGCTCTTCTCCGGAGACCTCTATAGTCTTCTAAATTTCCTCAGTTTTGCCAGGTGGCTTTTTATGGGGCTGGCAGTCGCAGGACTGATTTATCTTCGATACAAACGCCCAGATATGCATCGTCCTTTCAAGGTGCCTCTCTTCATCCCGGCACTATTTTCCTTCACCTGCCTCTTCATGGTTGTCCTCTCTCTTTACTCGGACCCATTCAGCACCGGGGTCGGTTTTCTTATCACCTTGACTGGGGTCCCTGCATATTATCTCTTCATTGTATGGGACAAGAAACCCAAGTGGTTCAGACGATTATCAGAC AGAATAACCAGAACATTACAGATTATACTAGAAGTTGTACCAGAAGACTCTAAAGAATTATGAGAATTCGGGG “. This sequence was then cloned in the pMIG vector using EcoR1 (RO101S, New England Biolabs) and BglII (RO143, New England Biolabs) restriction enzymes. The shRNA targeting mouse *Phgdh* were obtained using the oligonucleotide sequences forward 5’-GATCCCCGTGGAGAAGCAGAACTTGATTCAAGAGATCAAGTTCTGCTTCTCCACTTTTTA-3’, reverse 5’-AGCTTAAAAAGTGGAGAAGCAGAACTTGATCTCTTGAATCAAGTTCTGCTTCTCCACGGG-3 (sh*Phgdh* #1) and forward 5’-GATCCCCGGAGGAGCTGATAGCTGAATTCAAGAGATTCAGCTATCAGCTCCTCCTTTTTA-3’, reverse 5’-AGCTTAAAAAGGAGGAGCTGATAGCTGAATCTCTTGAATTCAGCTATCAGCTCCTCCGGG-3’ (sh*Phgdh* #2). As a control, shRNA targeting firefly luciferase were obtained using the oligonucleotides sequences forward 5’-GCGTTATTTATCGGAGTTG-3’ and reverse 5’-CAACTCCGATAAATAACGC-3’ (shSCR). 3 µg of corresponding forward and reverse oligonucleotides were annealed and cloned into the retroviral GFP-encoded pSUPER vector (VEC-PRT-0006, Oligoengine) linearized using HindIII (RO104S, New England Biolabs) and BglII restrictions enzymes according to the manufacture’s instruction.

Upon gel purification (Monarch DNA Gel Extraction Kit, T10205, New England Biolabs), ligation was performed using the T4 DNA ligase (MO202S, Biolabs) with a ratio vector:insert of 1:4. The generated vectors were amplified in MAX Efficiency Stbl2 competent cells (10268019, Thermo Fisher Scientific) with ampicillin selection, purified and sequenced.

### Generation of stable genetically modified Eµ-*Myc* cells

Self-inactivating retroviruses were generated following transient co-transfection of 293T cells (CRL-1573, ATCC) with 8.6 µg of one of the following vectors, pMIG empty vector (CTL), pMIG-*Asns*-v5 (ASNS OE), pMIG-*Phgdh*-v5 (PHGDH OE), or pMIG-*slc7a11* (SLC7A11 OE), pSUPER-shSCR or pSUPER-sh*Phgdh* (#equence #1 or #2), 3 µg of envelope plasmid phCMV-VSV-G and 8.6 µg of MLV-Gag-Pol, using the classical calcium phosphate method. After 15 hours, transfected 293T cells were incubated in Opti-MEM reduced serum medium (11058021, Thermo Fisher Scientific) for 32h. Cell supernatant was centrifuged 3000g at 4°C overnight to concentrate retroviral particles prior titration. Eµ-*Myc* cells were then transduced with the resulting retroviruses using a multiplicity of infection (MOI) of 10. 72 hours following transduction, GFP^+^ Eµ-*Myc* cells were sorted (SH800S Cell Sorter, Sony).

### Determination of glutamine and glutamate concentrations in medium

The native *E-Coli* L-asparaginase (ASNase, Kidrolase) was incubated in asparagine-containing medium in the absence of cells, at indicated concentrations and at 37°C, 5% CO_2_. The concentrations of glutamine and glutamate (mM) in the medium were electro-enzymatically determined overtime, using the YSI 2950 Biochemistry Analyzer (Yellow Springs Instruments).

### Western blot analysis

Eµ-*Myc* cells isolated *ex-vivo* from inguinal B-cell lymphomas or cultivated *in vitro* were washed in PBS and lysed in laemmli buffer to extract total proteins. Frozen axillary B-cell lymphomas were homogenized using a stainless-steel tissue grinder (1292, BioSpec Products) and proteins were extracted using the laemmli lysis buffer. After protein quantification (23225, Pierce BCA Protein Assay kit, Thermo Fisher Scientific), 15 to 30 µg of whole-cell protein lysates were separated on 8% to 15% SDS polyacrylamide gels and transferred onto polyvinylidene difluoride membranes (Millipore). Membranes were then blotted with antibodies against ASNS (14681-1-AP, Proteintech, RRID: AB_2060119), PHGDH (14719-1-AP, Proteintech, RRID: AB_2283938), PSAT1 (10501-1-AP, Proteintech, RRID: AB_2172597), PSPH (14513-1-AP, Proteintech, RRID: AB_2171464), NRF2 (12721, Cell Signaling, RRID: AB_2715528), GLUL (11037-2-AP, Proteintech, RRID: AB_2110650), GOT1 (14886-1-AP, Proteintech, RRID: AB_2113630), γH2AX (ab81299, Abcam, RRID: AB_1640564), ATF4 (11815, Cell Signaling, RRID: AB_2616025) for figures 2h and 4, ATF4 (81798-2-RR, Proteintech; RRID: AB_3670506) for figures 2d, 5, 7 and S7, xCT/SLC7A11 (98051, Cell Signaling, RRID: AB_2800296), Poly/Mono-ADP Ribose (89190, Cell Signaling); Phospho-ATR (Ser 428) (2853, Cell Signaling, RRID: AB_2290281), ATR (13934, Cell Signaling, RRID: AB_2798347), Phospho-Chk1 (Ser345) (2348, Cell Signaling, RRID: AB_331212), Chk1 (2360, Cell Signaling, RRID: AB_2080320), Phospho-RPA32 /RPA2 (Ser33) (10148, Cell Signaling, RRID: AB_3099645), RPA32/RPA2 (52448, Cell Signaling, RRID: AB_2750889), Phospho-KAP1 (Ser824) (A300-767A, Thermo Fisher Scientific, RRID: AB_669740), KAP1 (4123, Cell Signaling, RRID: 2256670); V5 (R96025, Life Technologies, RRID: AB_159313) and ERK2 (sc-1647, Santa Cruz, RRID: AB_627547) Immunoreactive bands were detected with anti-mouse (7076, Cell Signaling Technology, RRID: AB_330924) or anti-rabbit (7074S, Cell Signaling Technology, RRID: AB_2099233) IgG horseradish peroxidase (HRP)-linked antibodies. Immunoblots were visualized (FUJIFILM LAS4000) by chemiluminescence using Pierce ECL Western Blotting substrates (32106, Pierce ECL, Thermo Fisher Scientific). ERK2 was used as a loading control in all immunoblotting experiments. When indicated, quantification of the level of indicated proteins was performed using ImageJ software. Relative quantification was obtained by normalizing data to ERK2 levels and to indicated control conditions. Whole-cell lysate obtained from Eµ-*Myc* cells treated with 1µg/ml of etoposide (HY-13629, MCE) for 3 hours or stimulated with H_2_O_2_ (0.5 mM) for 5 minutes were used as a positive control for γH2AX expression and PARylation, respectively.

### PHGDH activity

For *in vitro* samples, live Eµ-*Myc* cells were lysed for 10 minutes on ice, in buffer A, containing 20 mM of Tris pH 7.4, 5 mM of EDTA pH 8.0, 150 mM of NaCl, 0.5 % of NP40 (UN3500, EUROMEDEX) and 0.01 % of cocktail of protease and phosphatase inhibitors (1861280, Thermo Fisher Scientific). For *ex-vivo* samples, frozen B-cell lymphomas were homogenized using a stainless-steel tissue grinder (1292, BioSpec Products) and proteins were extracted in buffer A. Then, 10 to 30 µg of proteins were incubated in PHGDH activity buffer containing 51.5 mM of Tris pH 8.0, 10 mM of MgCl_2_, 0.05 % of BSA (P06-1403500, Panbiotech), 0.01 % of Tween 20 (2001-B, Euromedex), 1.25 mM of L-glutamate (49449, Sigma), 0.3 mM of NAD^+^ (N1511, Sigma) and 0.1 mM of 3-phosphoglycerate (HY-141412, MedChemExpress) in a black 96-well plate (655079, Greiner). PHGDH activity was measured on a fluoroscan (Synergy H1, Biotek) at 460 nm after excitation at 355 nm as the increase in fluorescence related to NADH accumulation. PHGH activity is determined as the delta absorbance, representing the speed of NADH production. Data are normalized to milligrams of protein and expressed as mean ± SD of the percentage relative to the control condition (+Asn, *in vitro* or Vehicle, *in vivo*).

### L-[^3^H(G)]-Serine uptake

A total of 250.000 live Eµ-*Myc* cells/ml were seeded in asparagine, serine, glycine-free medium supplemented with 0.4 mM of L-serine (S4311, Sigma), 0.4 mM of L-glycine (G8790, Sigma), and with or without 0.37 mM of L-asparagine (A0884, Sigma) and 0.003 IU/ml of *E-coli* L-asparaginase (Kidrolase) for 24 hours at 37°C, 5% CO_2_. The number of live cells was determined by 4’,6-diamidino-2-phenylindole (DAPI, D9642, Sigma) (0.5 µg/ml) staining and flow cytometry analysis (Miltenyi Biotech). Then, 1x10^6^ of live Eμ-*Myc* cells (DAPI negative) were washed twice with PBS and incubated in a 12-well plate in 1mL of pre-warmed asparagine, serine, glycine-free medium supplemented as described above, with L-serine being replaced by 1 μCi/ml or 0,4 μCi/ml of L-[^3^H(G)]-Serine (NET248250UC, PerkinElmer), for 15 minutes or 3 hours at 37°C, 5% CO_2_, respectively. Subsequently, cells were washed twice with pre-chilled PBS and lysed with 120 μl of 0.1N NaOH and mixed with 4 ml of Ultima Gold (6013321, PerkinElmer). Radioactivity (count per minute, cpm) was measured using a β-scintillation counter (Tri-Carb 2810 TR, Perkin Elmer). L-[^3^H(G)]-serine transport rate was expressed as mean ± SD of cpm/10^6^ cells.

### Cell death assay

A total of 250.000 live Eµ-*Myc* cells/ml were treated with either DMSO (0.075%), or *E-Coli* L-asparaginase (0.003 IU/ml), or indicated concentration of PHGDH inhibitor BI-4916 (HY-126253, MCE), BI-4924 (HY-126254, MCE), NCT-503 (HY-101966, MCE), CBR-5884 (HY-100012, MCE), PKUMDL-WQ2101 (HY-123269, MCE), or Olaparib (AZD2281, HY-10162, MCE), or the combination of indicated drugs, in asparagine-free medium supplemented or not with 0.37 mM of L-asparagine (A0884, Sigma) at 37°C, 5% CO_2_ for the indicated time. For quenching ROS, 5 mM of N-acetyl-L-cysteine (A9165, Sigma) was added simultaneously. For inhibition of caspases, 20 µM of the pan-caspase inhibitor qVD-OPh (A1901, APExBIO) was added simultaneously. Following 24 or 48 hours of treatment, cells were labeled with 0.5 µg/ml of DAPI (D9642, Sigma) and immediately analyzed by flow cytometry (Miltenyi Biotec). Data represents the percentage of dead cells (DAPI+) and are expressed as mean ± SD.

### Cell proliferation assay

A total of 250.000 live Eµ-*Myc* cells/ml were treated with either DMSO (0.075%), or 0.003 IU/ml of *E-Coli* L-asparaginase, or 10 µM of BI-4916 (HY-126253, MCE), or 60 µM of BI-4924 (HY-126254, MCE), or 0.3 µM of Olaparib (AZD2281, HY-10162, MCE), or the combination of indicated drugs, in asparagine-free medium supplemented or not with 0.37 mM of L-asparagine, at 37°C, 5% CO2 for the indicated period.

For cell proliferation in asparagine, serine, glycine-free medium, a total of 250.000 live Eµ-*Myc* cells/ml were seeded in the presence or absence of L-asparagine (0.37 mM), and/or L-serine (0.4 mM)/L-glycine (0.4 mM) at 37°C, 5% CO2 for the indicated period.

Cells were diluted in the medium of the corresponding condition when they reached 1.10^6^ cells/ml. The number of live cells was determined overtime by DAPI (0.5 µg/ml) staining and flow cytometry analysis (Miltenyi Biotec). Data are expressed as the mean number of live cells (DAPI-) ± SD or ± SEM.

### Measurement of reactive oxygen species

A total of 250.000 live Eμ-*Myc* cells per ml were treated for 15 or 20 hours (as indicated in the legends) with either DMSO (0.075%), or 0.003 IU/ml of *E-Coli* L-asparaginase, or 10 µM of BI-4916 (HY-126253, MCE), or the combination of both drugs, in asparagine-free medium supplemented or not with L-asparagine (0.37 mM) at 37°C, 5% CO_2_. Cells were then counted by flow cytometry following DAPI staining (0.5 µg/ml) and 1x10^6^ live cells (DAPI negative) were seeded in 2 ml of their respective medium in 12-well plates and treated or not with 10 mM of N-acetyl-L-cysteine (A9165, Sigma) 1-hour prior addition of 2,5 μM of CellROX green probe (C10444, Invitrogen) for additional 30 minutes incubation at 37°C, 5% CO_2_. Cells were then collected and washed once with PBS, resuspended in 200uL of PBS, labelled with 0.5 µg/ml of DAPI (D9642, Sigma) and immediately analyzed by flow cytometry (Miltenyi Biotec). CellROX green is a cell-permeable and non-fluorescent reagent when in a reduced state. Upon oxidation, CellROX green exhibits a strong fluorogenic signal (excitation: 485 nm; emission: 520 nm) that is localized primarily in the nucleus and in the mitochondria, providing a reliable measure of reactive oxygen species (ROS) in live cells. Data are represented as the Median Fluorescence intensity (MFI) of the probe detected in live (DAPI-) cells, normalized to control condition (+Asn) and expressed as mean ± SD.

### 3D cell culture of HCT116 cells

HCT116 spheroids were prepared by seeding 3000 cells/well in 96-well ultra-low attachment plates (650970, Greiner Bio-One). After 72 hours, the indicated treatments were applied. Cell viability was assessed by adding propidium iodide (final concentration 0.3 mg/L) to each well at the endpoint. After a 30-minute incubation, brightfield and fluorescence images were captured using a fluorescence microscope. The images were then analyzed using the FIJI/IMAGEJ software (National Institutes of Health) by calculating the corrected total cell fluorescence (CTCF) using the formula CTCF = integrated density – (area of fluorescent object × mean fluorescence of background readings). The images were artificially colored and cropped for publication.

### *In vitro* stable isotope tracing

For α-15N-L-glutamine tracing experiment, 300.000 live Eµ-*Myc* cells/ml were incubated in asparagine-free medium supplemented with 2 mM of α-15N-L-glutamine (NLM-1016-PK, Cambridge Isotope Laboratories) instead of unlabeled L-glutamine, and with or without 0.37 mM of L-asparagine (A0884, Sigma), at 37°C, 5% CO2, for 24 hours.

For ^13^C2-Glycine tracing experiment, 300.000 live Eµ-*Myc* cells/ml were incubated in asparagine, serine, glycine-free medium supplemented with 0.4 mM of L-serine (S4311, Sigma), 0.4 mM of ^13^C2-Glycine (CLM-1017, Cambridge Isotope Laboratories), and with or without 0.37 mM of L-asparagine, at 37°C, 5% CO2, for 24 hours.

For [U-13C]-glucose tracing experiment, 300.000 live Eµ-*Myc* cells/ml were incubated in glucose and asparagine-free medium supplemented with 25 mM of [U-13C]-glucose (CML-1396, Cambridge Isotope Laboratories) and with or without L-asparagine (0.37 mM) and *E-Coli* L-asparaginase (0.003 IU/ml), at 37°C, 5% CO2, for 24 hours.

At endpoint, Eµ-*Myc* cells were analyzed by flow cytometry to determine the number of live cells (DAPI-) per mL. Then, 1·10^6^ live Eµ-*Myc* cells per replicate were collected, washed twice in pre-chilled PBS (300g, 5 min, 15°C centrifugation) and snap-frozen. Samples were stored at -80°C for further LC-MS-based metabolomic analyses. Results have been corrected for the presence of naturally occurring ^15^N or ^13^C stable isotopes using Metabolite AutoPlotter ^62^.

### *Ex-vivo* ATP analysis

ATP was measured using the Cell Titer Glo kit (G7570, Promega). Briefly, 20,000 malignant B cells harvested from freshly isolated B-cell lymphomas were resuspended in 80 µL of DMEM medium supplemented with 10% FBS, 2 mM of L-Glutamine and distributed in a 96 well-plate. Cells were then treated in triplicates for one hour with either control (DMSO), or 10 mg/ml of oligomycin (O4876, Sigma-Aldrich) or 10 mM iodoacetate (I9148, Sigma-Aldrich), or the combination of both metabolic inhibitors to obtain the residual amount of ATP. Short time treatment (1hour) was chosen to avoid cell death upon inhibition of both metabolic pathways. 100 µl of Cell Titer Glo reaction mix was then added in each well for a final volume of 200 µl. Plates were analyzed for luminescence with a Luminoscan (Berthold Technologies). We verified that ATP measurements were in the linear range of the detection. The difference between total ATP production and the ATP produced under oligomycin treatment results in OxPhos ATP contribution. OxPhos ATP is represented by percentage of total ATP produced by the cells.

### *In vivo* stable isotope tracing

When lymphoma development reached the ethical endpoint, mice received two consecutive intraperitoneal discrete boluses injections at 20 min interval of 1 g/kg of [U-^13^C_6_]-Glucose (CLM-1396, Cambridge Isotope Laboratories) in sterile NaCl 0.9% (100 mg/ml). Blood was collected from the tail-vein 40 min after the first [U-^13^C_6_]-Glucose bolus and plasma was isolated by centrifugation at 6000g for 2 min and snap-frozen. Mice were immediately sacrificed by cervical dislocation. All B-cell lymphomas were harvested and snap-frozen. All samples collected were stored at -80°C and axillary BCL were later used for LC-MS-based metabolomic, immunoblot and other indicated biochemical analyses. Results have been corrected for the presence of naturally occurring ^13^C stable isotopes using Metabolite AutoPlotter ^62^.

### Steady state metabolomics *in vitro*

250.000 live Eµ-*Myc* cells/ml were treated with indicated drugs for 24 hours, in asparagine-containing medium or in asparagine, serine, glycine-free medium supplemented with or without L-asparagine (0.37 mM), L-serine (0.4 mM) and glycine (0.4 mM) at 37°C, 5% CO_2_. The number of live cells was determined overtime by DAPI (0.5 µg/ml) staining and flow cytometry analysis (Miltenyi Biotec). Then, 1·10^6^ live cells (DAPI-) per replicate were collected, washed twice in pre-chilled PBS (300g, 5 min, 15°C centrifugation) and snap frozen. Samples were stored at -80°C for further LC-MS-based metabolomic analyses. The abundance of metabolite is represented as peak area. The GSSG/GSH ratio was calculated from total GSH and GSSG levels determined by LC-MS.

### Targeted LC/MS metabolites analyses

For *in vitro* samples, frozen pellets of 1·10^6^ Eµ-*Myc* cells were used for metabolite extraction. For lymphomas, frozen axillary B-cell lymphomas were homogenized using a stainless-steel tissue grinder (1292, BioSpec Products) and 10 mg of tissue were used for metabolite extraction. Extraction solution was composed of 50% methanol, 30% acetonitrile (ACN) and 20% water. The volume of the extraction solution was adjusted to the cell number or tissue weight (1 ml per 1million cells or 1 ml per 50 mg of tissues, respectively). Plasma was diluted 20 folds with the same extraction solvent. After addition of extraction solution, samples were vortexed for 5 min at 4 °C and centrifuged at 16,000*g* for 15 min at 4 °C. The supernatants were collected and stored at −80 °C until analysis. LC/MS analyses were conducted on a QExactive Plus Orbitrap mass spectrometer equipped with an Ion Max source and a HESI II probe coupled to a Dionex UltiMate 3000 UPLC system (Thermo Fisher Scientific). External mass calibration was performed using a standard calibration mixture every seven days, as recommended by the manufacturer. The 5 µl samples were injected onto a ZIC-pHILIC column (150 mm × 2.1 mm; i.d. 5 µm) with a guard column (20 mm × 2.1 mm; i.d. 5 µm) (Millipore) for LC separation. Buffer A was 20 mM ammonium carbonate, 0.1% ammonium hydroxide (pH 9.2), and buffer B was ACN. The chromatographic gradient was run at a flow rate of 0.200 µl min^−1^ as follows: 0–20 min, linear gradient from 80% to 20% of buffer B; 20–20.5 min, linear gradient from 20% to 80% of buffer B; 20.5–28 min, 80% buffer B. The mass spectrometer was operated in full scan, polarity switching mode with the spray voltage set to 2.5 kV and the heated capillary held at 320 °C. The sheath gas flow was set to 20 units, the auxiliary gas flow to 5 units and the sweep gas flow to 0 units. The metabolites were detected across a mass range of 75–1,000 *m*/*z* at a resolution of 35,000 (at 200 *m*/*z*) with the automatic gain control target at 10^6^ and the maximum injection time at 250 ms. Lock masses were used to ensure mass accuracy below 5 ppm. Data were acquired with Thermo Xcalibur software (Thermo). The peak areas of metabolites and isotopologues were determined using Thermo TraceFinder software (Thermo), identified by the exact mass of each singly charged ion and by the known retention time on the HPLC column. For the absolute quantification standard addition method was used^63^. For the Serine absolute quantification in plasma a standard addition method was used, where the addition of d-Serine to a pooled sample was carried out and the endogenous levels were subtracted from the spiked standards. Calibration curve was prepared in concentration range of 0.05-1.5 mM in a mixed sample to overcome possible matrix effects. Importantly, the preparation and processing of the calibration curve samples mirrored that of the experimental samples to ensure consistency.

## Data availability

For RNAseq analysis from Eµ-*Myc* cells, the data generated are available from the corresponding authors upon request. Metabolomics data are available in supplementary tables S1 and S2, related to figure 1. All other metabolomics data generated in this study are available from the corresponding authors upon request.

## Statistical analysis

Metabolites multivariate statistical analysis was performed with the Metaboanalyst 5.0 software (https://www.metaboanalyst.ca/). Metabolite relative concentration was normalized by median, and the data was auto-scaled (mean-centered and divided by the standard deviation of each variable) prior to analysis. After scaling the data, we performed principal component analysis. Univariate analysis was performed comparing metabolite levels between groups where metabolite differences were defined by the following fold change threshold 0.8 < Fold change (ASNase/Vehicle) >1.25 (which corresponds to log_2_ transformed threshold -0.3219 < log_2_ (Fold change (ASNase/Vehicle)) > + 0.3219) and significance as raw P-value < 0.05 (from non-parametric test), with unequal variance, and application of appropriated statistical test. Volcano plots and Metabolite Set Enrichment Analysis performed based on statistically significant discriminant metabolites, were built using the Metaboanalyst software and the Kyoto Encyclopedia of Genes and Genomes (KEGG) pathway computational tool.

All other statistical analyses and graphical representation of data was performed using GraphPad Prism 9 software. Comparative tests were considered significant if a 2-sided p <0.05. For *in vitro* experiments, results are expressed as mean ± SD (or ± SEM) as indicated in the figure legends. For *ex-vivo* experiments, results are expressed as mean ± SD. Statistical differences were determined by appropriated statistical test (two-sided Student’s t-test, one way or two-way ANOVA test, followed by multiple comparison test). A P-value < 0.05 was considered statistically significant.

## Acknowledgements

We gratefully acknowledge the Centre Méditerranéen de Médecine Moléculaire (C3M) animal facility. This work has been supported by la Ligue Nationale Contre le Cancer “Équipe Labellisée”, Institut National du Cancer (INCa PLBIO), la Fondation ARC pour la recherche sur le cancer, le Cancéropôle PACA and l’Agence Nationale de la Recherche (LABEX SIGNALIFE ANR-11-LABX-0028-01). Samples acquisition and data analysis were performed at C3M Cytometry Core Facility financed by Conseil Général CG06 and Conseil Régional PACA.

## Conflict of interest

The authors declare no conflict of interest.

## Legends to supplementary Figures

**Figure S1: ASNase treatment reprograms B-cell lymphomas metabolism *in vivo,* related to figure 1.**

**a.** Kaplan-Meier curves for the percentage of tumor-free mice following the intravenously injection of OxPhos-dependent Eμ-*Myc* #506 cells isolated from the Eμ-*Myc*^Tg/+^ mouse #506, into syngeneic wild-type C57BL/6 mice. Seven days post-tumor cell inoculation, mice were treated with Vehicle or *E-Coli* L-asparaginase (ASNase) every 48 hours until B-cell lymphoma (BCL) reached the ethical endpoint (n=25 Vehicle-treated mice; n=26 ASNase-treated mice; 3 experiments). P-value from log-rank test.

**b.** As in **a.** with OxPhos-dependent Eµ-*Myc* #688 cells isolated from the Eμ-*Myc*^Tg/+^ mouse #688 (n=12 mice/group; 2 experiments). P-value from log-rank test.

**c.** Plasma concentration of asparagine (Asn) and aspartate (Asp) in C57BL/6 mice bearing -Eµ-Myc-BCL as described in **a.** and **b.** (n=20 mice/group from 2 experiments; from Eµ-*Myc* #506 cells, n=10 mice/group; from Eµ-*Myc* #688 cells, n=10 mice/group). Data are expressed as mean ± SD. P-value from t-test.

**d.** Contribution of OxPhos metabolism to ATP production in BCL harvested from C57BL/6 mice bearing Eµ-*Myc* #506 cells (left panel, n=15 Vehicle-treated mice and n=17 ASNase-treated mice) or bearing Eµ-*Myc* #688 cells (right panel, n=14 mice/group), treated with Vehicle or ASNase as in **a.** Data are expressed as mean ± SD. P-value from t-test.

**e.** Volcano plot highlighting significantly deregulated metabolites (0.8 < Fold change (ASNase/Vehicle) >1.25 and raw P-value < 0.05) in Eµ-*Myc* #688 cells-derived BCL described in Figure 1d. (n=10 mice/group). Nucleotide precursors labelled in blue, proteinogenic amino acid labelled in pink.

**f.** Glutamine and glutamate concentrations (mM) overtime, in glutamine and Asn-containing medium, following ASNase supplementation at indicated concentrations (IU/ml). Data are expressed as mean ± SD (n=3 experiments).

**g.** Percentage of dead Eμ-*Myc* (#688) cells (DAPI+) cultivated for 24 hours in Asn-free medium supplemented with (+) or without (-) Asn (0.37 mM) and ASNase (0.003 IU/ml). Data are expressed as mean ± SD (n= 6 experiments). P-value from t-test.

**h.** Proliferation of Eµ-*Myc* (#688) cells cultivated as in **g.** until 96 hours. Data are expressed as the mean number of live cells (DAPI-) ± SEM (n= 6 experiments).

**i. j.** Relative abundance (peak area) of total intracellular serine (**i.**) and glycine (**j.**) in live Eµ-*Myc* (#688) cells cultivated for 24 hours under conditions described in **g.** Data are expressed as mean ± SD (n=4 biological replicates and two technical replicates). P-value from t-test.

**, p < 0.01; ***, p < 0.001; ****, p < 0.0001.

**Figure S2: Characterization of the mechanism leading to increased serine levels in ASNase-treated malignant B cells *in vitro*, related to figure 2**.

**a.** Three-days proliferation of Eµ-*Myc* (#688) cells cultivated in Asn-, serine- and glycine-free medium, supplemented (+; left panel) or not (-; right panel) with Asn (N, 0.37 mM) and with or without serine and glycine (S/G, 0.4 mM/0.4 mM). Data are expressed as the mean number of live cells (DAPI-) ± SD (n= 3 experiments).

**b.** Relative mRNA levels of *Phgdh, Psat1, Psph* and *Asn*, *Gls*, involved in *de novo* serine biosynthesis and asparagine-glutamine metabolism, respectively, in Eµ-*Myc* (#688) cells cultivated for 48 hours in Asn-free medium, supplemented (+) or not (-) with Asn (0.37 mM) and ASNase (0.003 IU/ml). Data are normalized to *Rplp0* and control condition (+ Asn) and represented as mean ± SD (n=3 experiments). P-value from t-test.

**c.** Relative quantification of ATF4 in Eµ-*Myc* #506 (square, n=2 independent experiments) and in Eµ-*Myc* #688 (triangle, n=2 independent experiments) cells, following 24 hours of incubation in glutamine (Gln) and Asn-free medium supplemented (+) or not (-) with Gln (2 mM), Asn (0.37 mM) or ASNase (0.003 IU/ml), as in figure 2d. Data are represented as mean ± SD. P-value from P-value from t-test.

**d.** Percentage of PHGDH activity in Eµ-*Myc* cells cultivated for 24 hours in Asn-free medium, supplemented (+) or not (-) with Asn (0.37 mM) and ASNase (0.003 IU/ml). Data are expressed as mean ± SD (n=5 experiments). P-value from t-test.

**e.** Schematic representation of *de novo* serine, glycine and asparagine biosynthesis from [U-13C]-glucose or from α-15N-L-Glutamine.

**f.** Relative abundance (peak area) of ^13^C-labelled asparagine isotopologues m+2 (upper panel) and m+3 (lower panel) in Eµ-*Myc* (#688) cells cultivated for 24 hours in Asn-free medium supplemented with [U-^13^C]-glucose and with (+) or without (-) Asn (0.37 mM) and ASNase (0.003 IU/ml). Data are expressed as mean ± SD (n=4 biological replicates and two technical replicates). P-value from t-test.

**g.** Relative abundance (peak area) of Gln α-^15^N-labelled isotopologues in Eµ-*Myc* (#688) cells cultivated for 24 hours in Gln and Asn free-medium containing serine (0.4 mM) and glycine (0.4 mM), supplemented with α-^15^N-L-glutamine (2 mM) and with (+) or without (-) Asn (0.37 mM). Data are expressed as mean ± SD (n=4 biological replicates and two technical replicates).

**h.** Relative abundance (peak area) of α-^15^N-serine m+1 (left panel) and α-^15^N-Asn m+1 (right panel) in live Eµ-*Myc* (#688) cells cultivated as in **f.** Data are expressed as mean ± SD (n=4 biological replicates and two technical replicates). P-value from t-test.

**i.** Relative quantification of the expression level of PHGDH, PSAT1 and PSPH proteins in BCL harvested from Vehicle- and ASNase-treated mice bearing Eµ-*Myc* #506 cells, presented in figure 2h. (Vehicle, n=4 mice; ASNase, n=10 mice). Data are normalized to the Vehicle-treated condition. P-value from t-test.

*ns*, not significant; *, p < 0.05; **, p < 0.01; ***, p < 0.001; ****, p < 0.0001.

**Figure S3: *In vivo* [U-13C]-glucose tracing in Eµ-*Myc*-cells-bearing mice treated with Vehicle or ASNase, related to figure 2.**

**a.** Relative abundance (peak area) of ^13^C-labelled glucose isotopologues in the plasma of B-cell lymphomas (BCL)-bearing C57BL/6 mice that have been treated with Vehicle or *E-Coli* L-asparaginase (ASNase) every 48 hours until endpoint, prior two consecutive boluses of [U-^13^C]-glucose. Data are expressed as mean ± SD (n=4 mice/group). P-value from t-test.

**b. c. d.** Relative abundance (peak area) of serine (**b.**), glycine (**c.**) and asparagine (**d.**) in BCL harvested from Vehicle and ASNase-treated mice presented in **a.** Data are expressed as mean ± SD (n=4 mice/group). P-value from by t-test.

**e.** Relative abundance (peak area) of ^13^C-labelled asparagine m+2 isotopologue in BCL, following [U-^13^C]-glucose boluses in Eµ-*Myc* cells-bearing mice that have been treated with Vehicle or ASNase every 48 hours until endpoint. Data are expressed as mean ± SD (n=4 mice/group). P-value was determined by t-test.

**f.** Serine concentration (mM) in the plasma of Eµ-*Myc* cells-bearing C57BL/6 mice treated with Vehicle or ASNase every 48 hours until endpoint. Left panel: from mice injected with Eµ-*Myc* #506 cells, n=10 mice/group; right panel: from mice injected with Eµ-*Myc* #688 cells, n=16 mice/group). Data are expressed as mean ± SD. P-value from t-test.

*ns*, not significant; *, p < 0.05; **, p < 0.01; ***, p < 0.001; ****, p < 0.0001.

**Figure S4: ASNase-treated Eµ-*Myc* cells resume proliferation in a PHGDH - dependent manner, related to figure 3.**

**a. b. c. d.** Percentage of dead Eµ-*Myc* (#688) cells (DAPI+) following 24 hours of treatment with DMSO or indicated concentrations of the PHGDH inhibitors NCT-503 (**a.**), CBR-5884 (**b.**), PKUMDL-WQ2101 (**c.**), BI-4916 (**d.**) in Asn-containing medium. Data are expressed as mean ± SD (n= 3 experiments; **d.** n ≥5 experiments/condition). P-value from one-way Anova, followed by Tukey’s test.

**e.** Percentage of PHGDH activity in Eµ-*Myc* (#688) cells following 24 hours of treatment with indicated concentration of PHGDH inhibitors. Data are expressed as mean ± SD (n= 3 experiments). P-value from one-way Anova, followed by Tukey’s test.

**f.** Total serine levels in Eµ-*Myc* (#688) cells treated with DMSO, *E-Coli* L-asparaginase (ASNase, 0.003 IU/ml) and/or BI-4916 (10 µM) for 24 hours in Asn-containing medium). Data are expressed as mean ± SD (n=6 biological replicates). P-value from 2way Anova, followed by Tukey’s test.

**g.** Percentage of dead Eµ-*Myc* (#506) cells (DAPI+) following 24 hours of treatment with DMSO, *E-Coli* L-asparaginase (ASNase, 0.003 IU/ml) and/or BI-4916 (10 µM) in Asn-containing medium. Data are expressed as mean ± SD (n= 5 experiments). P-value from 2way Anova, followed by Tukey’s test.

**h.** As in **g.** in the presence or in the absence of the caspase inhibitor qVD-OPh (20 µM). Data are expressed as mean ± SD (n=3 independent experiments). P-value from 2way Anova, followed by Sidak’s test.

**i.** Four-days proliferation of Eµ-*Myc* (#506) cells treated as described in **g**. Data are expressed as the mean number of live cells (DAPI-) ± SD (n= 5 experiments). P-value from 2way Anova, followed by Tukey’s test.

**j.** PHGDH activity in Eµ-*Myc* (#506 and #688) cells treated with DMSO or indicated concentrations of the PHGDH inhibitor, BI-4924 or its pro-drug BI-4916 for 24 hours in Asn-containing medium (n=2 experiments with Eµ-*Myc* #688 cells and n=1 experiment with Eµ-*Myc* #506 cells). P-value from one-way Anova, followed by Tukey’s test.

**k.** Four-days proliferation of Eµ-*Myc* (#506) cells treated with DMSO, ASNase (0.003 IU/ml) and/or the PHGDH inhibitor BI-4924 (60 µM) in Asn-containing medium. Data are expressed as the mean number of live cells (DAPI-) ± SEM (n= 3 experiments). P-value from 2way Anova, followed by Tukey’s test.

**l.** Total protein extracts prepared from whole Eµ-*Myc* cells (#506) stably overexpressing control vector (CTL) or V5-tagged murine PHGDH (PHGDH OE) were immunoblotted for the indicated proteins. ERK2, loading control. Immunoblots shown are representative of 3 independent experiments.

**m.** Quantification of PHGDH, PSAT1 and PSPH expression in Eµ-*Myc* cells presented in **l.** (n=3 independent experiments). P-value from t-test.

**n.** Survival curves of WT C57BL/6 mice intravenously injected with Eµ-*Myc* (#506) cells stably overexpressing either control vector (CTL) or V5-tagged murine PHGDH (PHGDH OE) and treated 7 days later with either Vehicle or ASNase every 48 hours until endpoint (CTL-Vehicle, n=10 mice; CTL-ASNase, n=9 mice; PHGDH OE-Vehicle, n=10 mice; PHGDH OE-ASNase, n=10 mice). P-value from log-rank test.

*ns*, not significant; *, p < 0.05; **, p < 0.01; ***, p < 0.001; ****, p < 0.0001.

**Figure S5: Characterization of Eµ-*Myc* cell viability, proliferation *in vitro* and deregulated metabolic pathways *in vivo*, following a second ASNase therapy challenge, related to figure 4.**

**a.** Percentage of dead (DAPI+) WT and C1 cells derived Eµ-*Myc* #688 cells (as presented in figure 4a) treated or not (Ctl) with *E-Coli* L-asparaginase (ASNase, 0.003 IU/ml) in Asn-containing medium for 24 hours Data are expressed as mean ± SD (n= 3 experiments). P-value from t-test.

**b.** Proliferation of WT and C1 cells (derived from Eµ-*Myc* #688 cells) treated under conditions described in **a.** until 72 hours. Data are expressed as the mean number of live cells (DAPI-) ± SD (n= 3 experiments). P-value from t-test.

**c.** Overview of the top 18 significant enriched metabolite sets (KEGG library) from 78 significant deregulated metabolites (Fold change Vehicle/ASNase >1.25 and raw P-value < 0.05, from non-parametric test) in axillary B-cell lymphomas (BCL) (obtained following the intravenous injection of ^A^BCL cells into WT C57BL/6 mice and a 1^st^ ASNase therapy) following a second ASNase challenge *in vivo*.

**d.** Relative quantification of the expression levels of indicated proteins in BCL presented in figure 4j. Data are normalized to the ^A^BCL condition (resulting from the 1^st^ ASNase challenge) and expressed as mean ± SD (n=6 Vehicle ^A^BCL; n=6 ASNase-^V^BCL). P-value from t-test.

**Figure S6: Increased glutathione biosynthesis in asparagine-restricted Eµ-*Myc* cells and expression of *slc7a11* mRNA in malignant cells, related to figure 5.**

**a.** GSSG/GSH ratio in Eµ-*Myc* (#688) cells treated with DMSO or *E-Coli* L-asparaginase (ASNase, 0.003 IU/ml) and/or BI-4916 (10 µM) for 24 hours in Asn-containing medium. Data are expressed as mean ± SD (n=6 biological replicates). P-value from 2-way Anova followed by Tukey’s test.

**b.** Relative mRNA levels of the glycine transporter *slc6a9,* in Eµ-*Myc* cells cultivated in Asn-free medium, supplemented (+) or not (-) with Asn (0.37 mM) and ASNase (0.003 IU/ml) for 48 hours. Data are normalized to *Rplp0* and control condition (+ Asn) and represented as mean ± SD (n=3 experiments). P-value from t-test.

**c.** Relative abundance (peak area) of glycine isotopologues (left panel) and of carbon-labelled glycine m+2 (right panel) in Eµ-*Myc* (#688) cells following 24 hours incubation in Asn-, serine- and glycine-free medium supplemented with serine (0.4 mM), ^13^C_2_-glycine (0.4 mM), and with (+) or without Asn (0.37 mM).

**d.** As in **c.** for GSH isotopologues (left panel) and carbon labelled GSH m+2 (right panel).

**e.** Relative abundance (peak area) of other glycine-derived metabolites in Eµ-*Myc* cells (#688) cultivated as in **c.** From **c.** to **e.** data are represented as mean ± SD (n=4 biological replicates and two technical replicates). P value from t-test.

**f.** Relative levels of *slc7a11* mRNA in murine solid cancer cell lines (colorectal carcinoma, CT26, MC38; lung carcinoma, LLC1) and in Eµ-*Myc* (#688) cells cultivated for the indicated period in Asn-free medium, supplemented (+) or not (-) with Asn (0.37 mM) and ASNase (0.003 IU/ml). Data normalized to *Rplp0* and to CT26 cell line.

**g.** Heatmap representation of cystine transporters (*slc7a11, slc7a9, slc3a1*) mRNA expression (log_2_-transformed) in 756 human DLBCL, based on RNAseq data from ^42^

**|h.** 756 DLBCL samples were classified according to *slc7a11* mRNA expression (null/very low, 0≤log_2_< 3.0, n=55; 3≤log_2_<6; low, 6≤log_2_<10, n=216; intermediate 10≤log_2_<12, n=454; high, log_2_≥12, n=31) from RNAseq ^42^. The percentage of samples is reported above boxplot.

*ns*, not significant; *, p < 0.05; **, p < 0.01; ***, p < 0.001; ****, p < 0.0001.

**Figure S7: *In vivo* increase in γH2AX expression and the impact of PHGDH overexpression in B-cell lymphomas treated *in vivo* with ASNase, related to figure 6.**

**a.** Expression of indicated proteins in B-cell lymphomas (BCL) from Eµ-*Myc* (#506) cells-bearing C57BL/6 mice treated *in vivo* with either Vehicle (NaCl 0.9%) or ASNase until endpoint. ERK2, loading control (n=7 mice/group).

**b. c.** Relative quantification of γH2AX expression (**b.**) and P-Chk1 (Ser345) (**c.**) in BCL presented in **a.** P-value from t-test.

*ns*, not significant; *, p < 0.05; ****, p < 0.0001.

**Figure S8: Combining ASNase and Olaparib is effective in Eµ-*Myc* cells and in a 3D cell culture model of HCT116 colon carcinoma cell line, related to figure 7.**

**a.** HCT116 colorectal cancer cells were cultured in ultra-low attachment conditions for 3 days, followed by treatment with DMSO, *E-Coli* L-asparaginase (ASNase, 0.2 U/mL), Olaparib (Ola, 0.1 μM), or the combination of the two drugs for five days. Cell death was assessed by propidium iodide (PI) incorporation, and spheroids were imaged.

**b.** Quantification PI incorporation in spheroids treated with DMSO, ASNase, Olaparib or ASNase/Olaparib combination as shown in **a.** Data are represented as mean ± SD (n=3 experiments). P-value from 2way Anova, followed by Tukey’s test.

*ns*, not significant; *, p < 0.05; ***, p < 0.001; ****, p < 0.0001.

**Table S1. Statistically significant discriminant metabolites identified in B-cell lymphomas following *in vivo* treatment with Vehicle or ASNase, related to figure 1**. Comparative metabolomic analysis of B-cell lymphomas (BCL) derived from Eµ-*Myc* #506 cells or Eµ-*Myc* #688 cells following *in vivo* treatment with either Vehicle or ASNase until lymphoma development reached the endpoint. Metabolites were considered significantly deregulated based on a fold change threshold of ASNase/Vehicle >1.25 and a raw P-value < 0.05 (non-parametric test). Across both #506-derived and #688-derived BCL datasets, a total of 14 metabolites were consistently increased (highlighted in red), and 5 were decreased (highlighted in blue) in response to ASNase treatment. Notably, 3 metabolites (highlighted in grey) displayed discordant regulation between #506-BCL and #688-BCL, suggesting potential clone-specific metabolic adaptation.

**Table S2: Fold changes in proteinogenic amino acids levels in B-cell lymphomas treated *in vivo* with ASNase *vs* Vehicle, related to figure 1**. Tables summarize the fold change and statistical significance in the abundance of proteinogenic amino acids (AAs) in B-cell lymphomas (BCL) derived from Eµ*-Myc* #506 cells (upper table) and Eµ-*Myc* #688 cells (lower table) following *in vivo* treatment with ASNase compared to Vehicle-treated controls. Metabolite abundance changes are reported as fold change of ASNase/Vehicle >1.25. Statistical significance was determined using a non-parametric test (raw P-value < 0.05) and all significantly deregulated AAs are indicated in bold. Color-coded highlights indicate the following: Bold black, statistically significant AAs that are either not shared between the two datasets, not consistently regulated in the same direction, or exhibit a fold change below the threshold; bold purple, the only AA that is significantly deregulated in both datasets but display a fold change < 1.25 in the #506-BCL dataset; bold blue, the four AAs that are significantly deregulated with a fold change ASNase/Vehicle>1.25, and consistently regulated in both #506-BCL and #688-BCL following ASNase treatment.

## Notes

### Competing Interest Statement

The authors have declared no competing interest.

## References

1. Schmidt, D.R. et al. Metabolomics in cancer research and emerging applications in clinical oncology. CA Cancer J Clin 71, 333–358 (2021).

2. Lemberg, K.M., Gori, S.S., Tsukamoto, T., Rais, R. & Slusher, B.S. Clinical development of metabolic inhibitors for oncology. J Clin Invest 132 (2022).

3. Stine, Z.E., Schug, Z.T., Salvino, J.M. & Dang, C.V. Targeting cancer metabolism in the era of precision oncology. Nat Rev Drug Discov 21, 141–162 (2022).

4. Vasan, K. & Chandel, N.S. Molecular and cellular mechanisms underlying the failure of mitochondrial metabolism drugs in cancer clinical trials. J Clin Invest 134 (2024).

5. Akagi, T. et al. Methylation analysis of asparagine synthetase gene in acute lymphoblastic leukemia cells. Leukemia 20, 1303–1306 (2006).

6. Touzart, A. et al. Epigenetic Silencing Affects l-Asparaginase Sensitivity and Predicts Outcome in T-ALL. Clin Cancer Res 25, 2483–2493 (2019).

7. Kucuk, C. et al. Global promoter methylation analysis reveals novel candidate tumor suppressor genes in natural killer cell lymphoma. Clin Cancer Res 21, 1699–1711 (2015).

8. Chiche, J. et al. GAPDH Expression Predicts the Response to R-CHOP, the Tumor Metabolic Status, and the Response of DLBCL Patients to Metabolic Inhibitors. Cell Metab 29, 1243–1257 e1210 (2019).

9. Grima-Reyes, M. et al. Tumoral microenvironment prevents de novo asparagine biosynthesis in B cell lymphoma, regardless of ASNS expression. Sci Adv 8, eabn6491 (2022).

10. Pathria, G. et al. Translational reprogramming marks adaptation to asparagine restriction in cancer. Nat Cell Biol 21, 1590–1603 (2019).

11. Gwinn, D.M. et al. Oncogenic KRAS Regulates Amino Acid Homeostasis and Asparagine Biosynthesis via ATF4 and Alters Sensitivity to L-Asparaginase. Cancer Cell 33, 91–107 e106 (2018).

12. Nakamura, A. et al. Inhibition of GCN2 sensitizes ASNS-low cancer cells to asparaginase by disrupting the amino acid response. Proc Natl Acad Sci U S A 115, E7776–E7785 (2018).

13. Cecconello, D.K. et al. Asparaginase: an old drug with new questions. Hematol Transfus Cell Ther 42, 275–282 (2020).

14. Mondelaers, V. et al. Prospective, real-time monitoring of pegylated Escherichia coli and Erwinia asparaginase therapy in childhood acute lymphoblastic leukaemia and non-Hodgkin lymphoma in Belgium. Br J Haematol 190, 105–114 (2020).

15. Schore, R.J. et al. Plasma asparaginase activity and asparagine depletion in acute lymphoblastic leukemia patients treated with pegaspargase on Children’s Oncology Group AALL07P4(). Leuk Lymphoma 60, 1740–1748 (2019).

16. Iwamoto, S., Mihara, K., Downing, J.R., Pui, C.H. & Campana, D. Mesenchymal cells regulate the response of acute lymphoblastic leukemia cells to asparaginase. J Clin Invest 117, 1049–1057 (2007).

17. Ehsanipour, E.A. et al. Adipocytes cause leukemia cell resistance to L-asparaginase via release of glutamine. Cancer Res 73, 2998–3006 (2013).

18. Tardito, S. et al. The inhibition of glutamine synthetase sensitizes human sarcoma cells to L-asparaginase. Cancer Chemother Pharmacol 60, 751–758 (2007).

19. Rotoli, B.M. et al. Inhibition of glutamine synthetase triggers apoptosis in asparaginase-resistant cells. Cell Physiol Biochem 15, 281–292 (2005).

20. Guarecuco, R. et al. Dietary thiamine influences l-asparaginase sensitivity in a subset of leukemia cells. Sci Adv 6 (2020).

21. Krall, A.S. et al. Asparagine couples mitochondrial respiration to ATF4 activity and tumor growth. Cell Metab 33, 1013–1026 e1016 (2021).

22. Farge, T. et al. Chemotherapy-Resistant Human Acute Myeloid Leukemia Cells Are Not Enriched for Leukemic Stem Cells but Require Oxidative Metabolism. Cancer Discov 7, 716–735 (2017).

23. Denise, C. et al. 5-fluorouracil resistant colon cancer cells are addicted to OXPHOS to survive and enhance stem-like traits. Oncotarget 6, 41706–41721 (2015).

24. Ippolito, L. et al. Metabolic shift toward oxidative phosphorylation in docetaxel resistant prostate cancer cells. Oncotarget 7, 61890–61904 (2016).

25. Krall, A.S., Xu, S., Graeber, T.G., Braas, D. & Christofk, H.R. Asparagine promotes cancer cell proliferation through use as an amino acid exchange factor. Nat Commun 7, 11457 (2016).

26. Chen, H., Pan, Y.X., Dudenhausen, E.E. & Kilberg, M.S. Amino acid deprivation induces the transcription rate of the human asparagine synthetase gene through a timed program of expression and promoter binding of nutrient-responsive basic region/leucine zipper transcription factors as well as localized histone acetylation. J Biol Chem 279, 50829–50839 (2004).

27. Gao, S. et al. PSAT1 is regulated by ATF4 and enhances cell proliferation via the GSK3beta/beta-catenin/cyclin D1 signaling pathway in ER-negative breast cancer. J Exp Clin Cancer Res 36, 179 (2017).

28. Riscal, R. et al. Chromatin-Bound MDM2 Regulates Serine Metabolism and Redox Homeostasis Independently of p53. Mol Cell 62, 890–902 (2016).

29. Tameire, F. et al. ATF4 couples MYC-dependent translational activity to bioenergetic demands during tumour progression. Nat Cell Biol 21, 889–899 (2019).

30. Bollino, D. et al. Erwinia asparaginase (crisantaspase) increases plasma levels of serine and glycine. Front Oncol 12, 1035537 (2022).

31. Pacold, M.E. et al. A PHGDH inhibitor reveals coordination of serine synthesis and one-carbon unit fate. Nat Chem Biol 12, 452–458 (2016).

32. Mullarky, E. et al. Identification of a small molecule inhibitor of 3-phosphoglycerate dehydrogenase to target serine biosynthesis in cancers. Proc Natl Acad Sci U S A 113, 1778–1783 (2016).

33. Wang, Q. et al. Rational Design of Selective Allosteric Inhibitors of PHGDH and Serine Synthesis with Anti-tumor Activity. Cell Chem Biol 24, 55–65 (2017).

34. Weinstabl, H. et al. Intracellular Trapping of the Selective Phosphoglycerate Dehydrogenase (PHGDH) Inhibitor BI-4924 Disrupts Serine Biosynthesis. J Med Chem 62, 7976–7997 (2019).

35. Kiweler, N. et al. Mitochondria preserve an autarkic one-carbon cycle to confer growth-independent cancer cell migration and metastasis. Nat Commun 13, 2699 (2022).

36. Graham, N.A. et al. Glucose deprivation activates a metabolic and signaling amplification loop leading to cell death. Mol Syst Biol 8, 589 (2012).

37. Hamano, M. et al. Enhanced vulnerability to oxidative stress and induction of inflammatory gene expression in 3-phosphoglycerate dehydrogenase-deficient fibroblasts. FEBS Open Bio 8, 914–922 (2018).

38. Zgheib, E. et al. Investigation of Nrf2, AhR and ATF4 Activation in Toxicogenomic Databases. Front Genet 9, 429 (2018).

39. Hayes, J.D., Dinkova-Kostova, A.T. & Tew, K.D. Oxidative Stress in Cancer. Cancer Cell 38, 167–197 (2020).

40. Cheung, E.C. & Vousden, K.H. The role of ROS in tumour development and progression. Nat Rev Cancer 22, 280–297 (2022).

41. Bansal, A. & Simon, M.C. Glutathione metabolism in cancer progression and treatment resistance. J Cell Biol 217, 2291–2298 (2018).

42. Reddy, A. et al. Genetic and Functional Drivers of Diffuse Large B Cell Lymphoma. Cell 171, 481–494 e415 (2017).

43. Sun, J. et al. SLC1A3 contributes to L-asparaginase resistance in solid tumors. EMBO J 38, e102147 (2019).

44. Hinze, L. et al. Synthetic Lethality of Wnt Pathway Activation and Asparaginase in Drug-Resistant Acute Leukemias. Cancer Cell 35, 664–676 e667 (2019).

45. Apfel, V. et al. Therapeutic Assessment of Targeting ASNS Combined with l-Asparaginase Treatment in Solid Tumors and Investigation of Resistance Mechanisms. ACS Pharmacol Transl Sci 4, 327–337 (2021).

46. Butler, M. et al. BTK inhibition sensitizes acute lymphoblastic leukemia to asparaginase by suppressing the amino acid response pathway. Blood 138, 2383–2395 (2021).

47. D’Avola, A. et al. PHGDH is required for germinal center formation and is a therapeutic target in MYC-driven lymphoma. J Clin Invest 132 (2022).

48. Hope, H.C., et al. Coordination of asparagine uptake and asparagine synthetase expression modulates CD8+ T cell activation. JCI Insight 6 (2021).

49. Wu, J. et al. Asparagine enhances LCK signalling to potentiate CD8(+) T-cell activation and anti-tumour responses. Nat Cell Biol 23, 75–86 (2021).

50. Gnanaprakasam, J.N.R. et al. Asparagine restriction enhances CD8(+) T cell metabolic fitness and antitumoral functionality through an NRF2-dependent stress response. Nat Metab 5, 1423–1439 (2023).

51. Chang, H.C. et al. Asparagine deprivation enhances T cell antitumour response in patients via ROS-mediated metabolic and signal adaptations. Nat Metab 7, 918–927 (2025).

52. Strom, C.E. et al. Poly (ADP-ribose) polymerase (PARP) is not involved in base excision repair but PARP inhibition traps a single-strand intermediate. Nucleic Acids Res 39, 3166–3175 (2011).

53. Galindo-Campos, M.A. et al. Distinct roles for PARP-1 and PARP-2 in c-Myc-driven B-cell lymphoma in mice. Blood 139, 228–239 (2022).

54. Elfar, G.A. et al. p53-dependent crosstalk between DNA replication integrity and redox metabolism mediated through a NRF2-PARP1 axis. Nucleic Acids Res 52, 12351–12377 (2024).

55. Ferguson, D.C. et al. Amino acid stress response genes promote L-asparaginase resistance in pediatric acute lymphoblastic leukemia. Blood Adv 6, 3386–3397 (2022).

56. Bryant, H.E. et al. Specific killing of BRCA2-deficient tumours with inhibitors of poly(ADP-ribose) polymerase. Nature 434, 913–917 (2005).

57. Farmer, H. et al. Targeting the DNA repair defect in BRCA mutant cells as a therapeutic strategy. Nature 434, 917–921 (2005).

58. Parvin, S. et al. LMO2 Confers Synthetic Lethality to PARP Inhibition in DLBCL. Cancer Cell 36, 237–249 e236 (2019).

59. Ricci, A.D. et al. Specific Toxicity of Maintenance Olaparib Versus Placebo in Advanced Malignancies: A Systematic Review and Meta-analysis. Anticancer Res 40, 597–608 (2020).

60. Rose, M., Burgess, J.T., O’Byrne, K., Richard, D.J. & Bolderson, E. PARP Inhibitors: Clinical Relevance, Mechanisms of Action and Tumor Resistance. Front Cell Dev Biol 8, 564601 (2020).

61. Chiche, J. et al. GAPDH enhances the aggressiveness and the vascularization of non-Hodgkin’s B lymphomas via NF-kappaB-dependent induction of HIF-1alpha. Leukemia 29, 1163–1176 (2015).

62. Pietzke, M. & Vazquez, A. Metabolite AutoPlotter - an application to process and visualise metabolite data in the web browser. Cancer Metab 8, 15 (2020).

63. Luo, B., Groenke, K., Takors, R., Wandrey, C. & Oldiges, M. Simultaneous determination of multiple intracellular metabolites in glycolysis, pentose phosphate pathway and tricarboxylic acid cycle by liquid chromatography-mass spectrometry. J Chromatogr A 1147, 153–164 (2007).

