## Supplementary figures for "L-asparaginase treatment induces a tumor metabolic plasticity and reveals a vulnerability to PARP1/2 inhibitor Olaparib, in B-cell lymphomas"

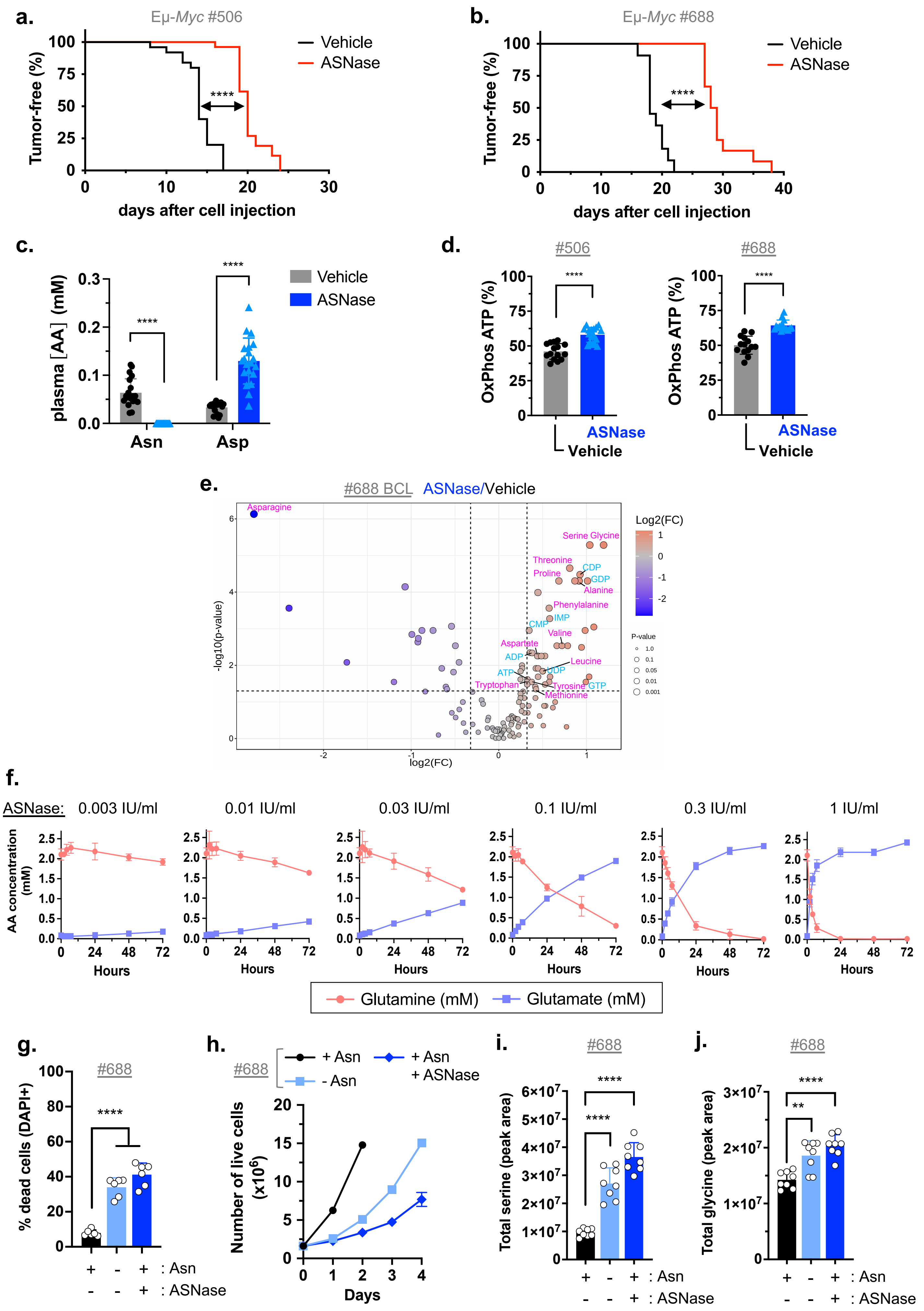


Aussel A. *et al.* Figure S1

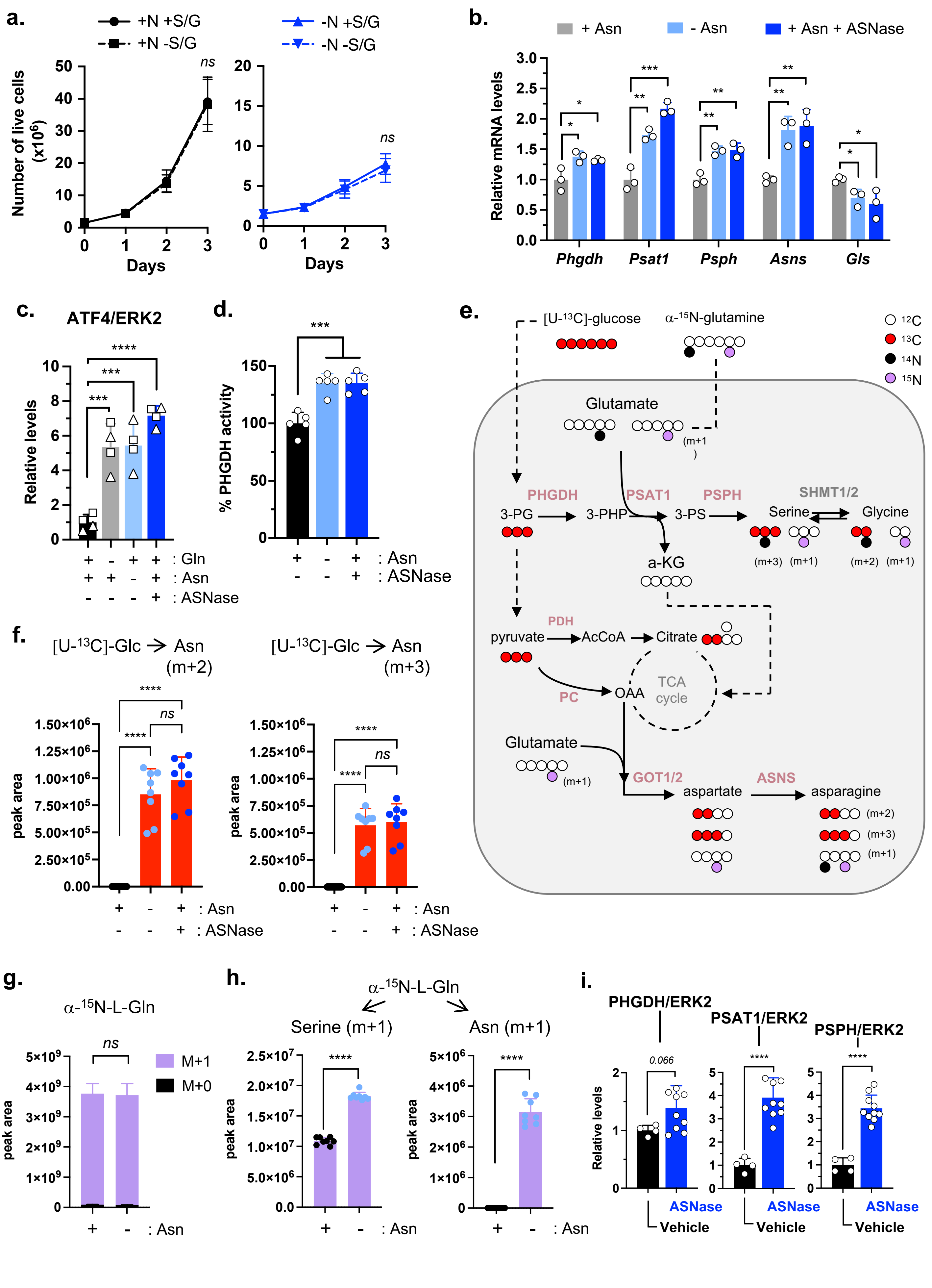


Aussel A. *et al.* Figure S2

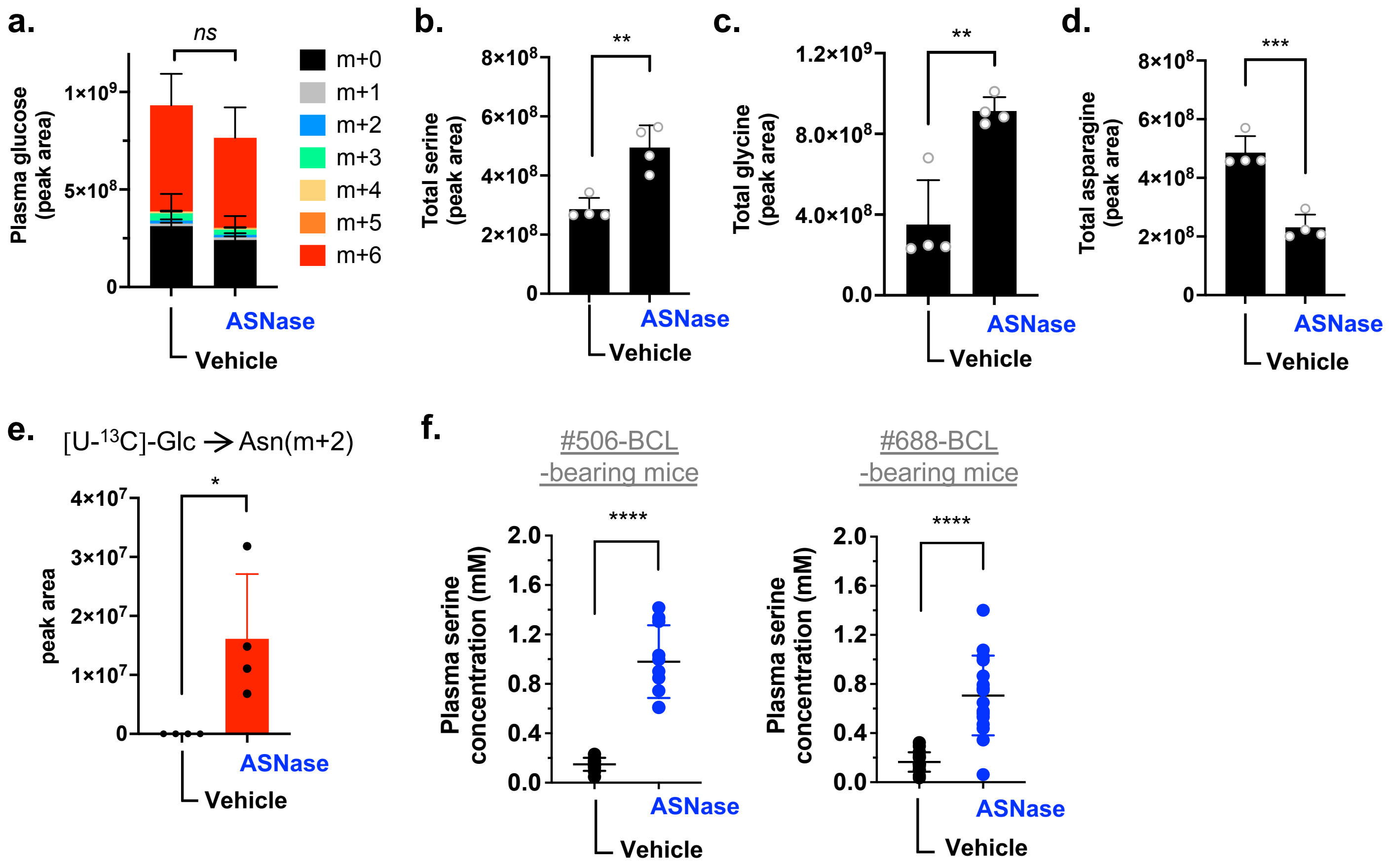


Aussel A. *et al.* Figure S3

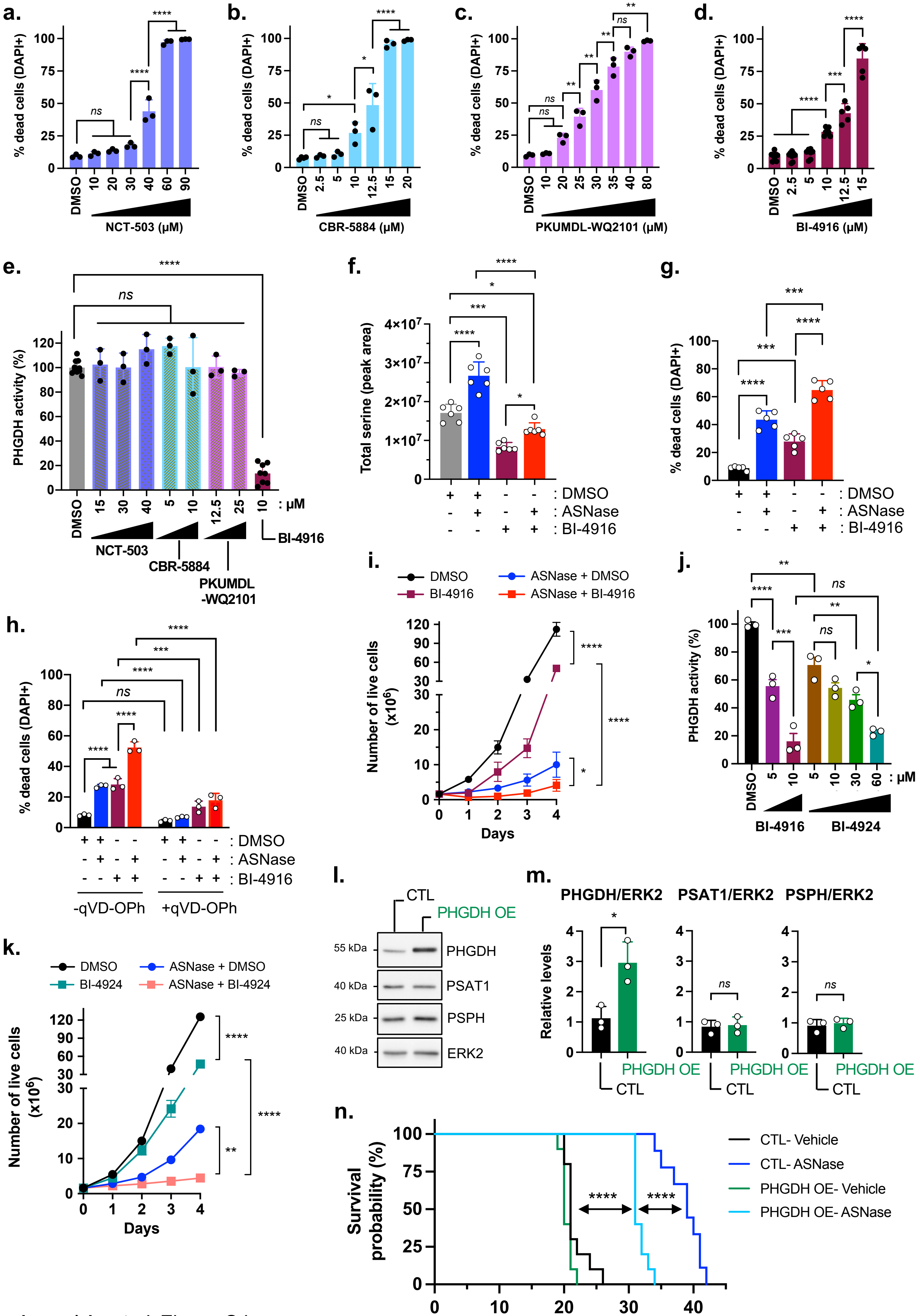


Aussel A. *et al.* Figure S4

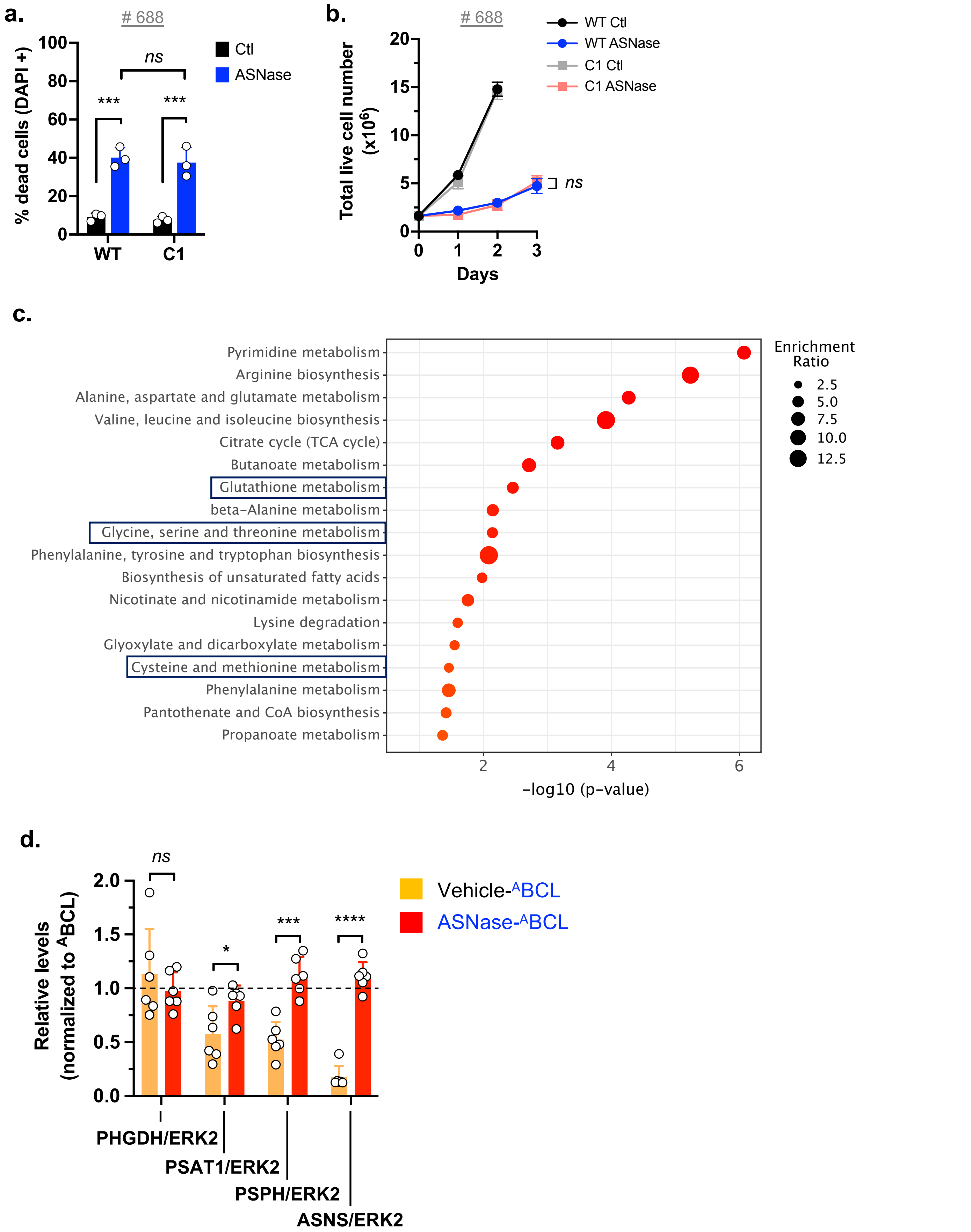


Aussel A. *et al.* Figure S5

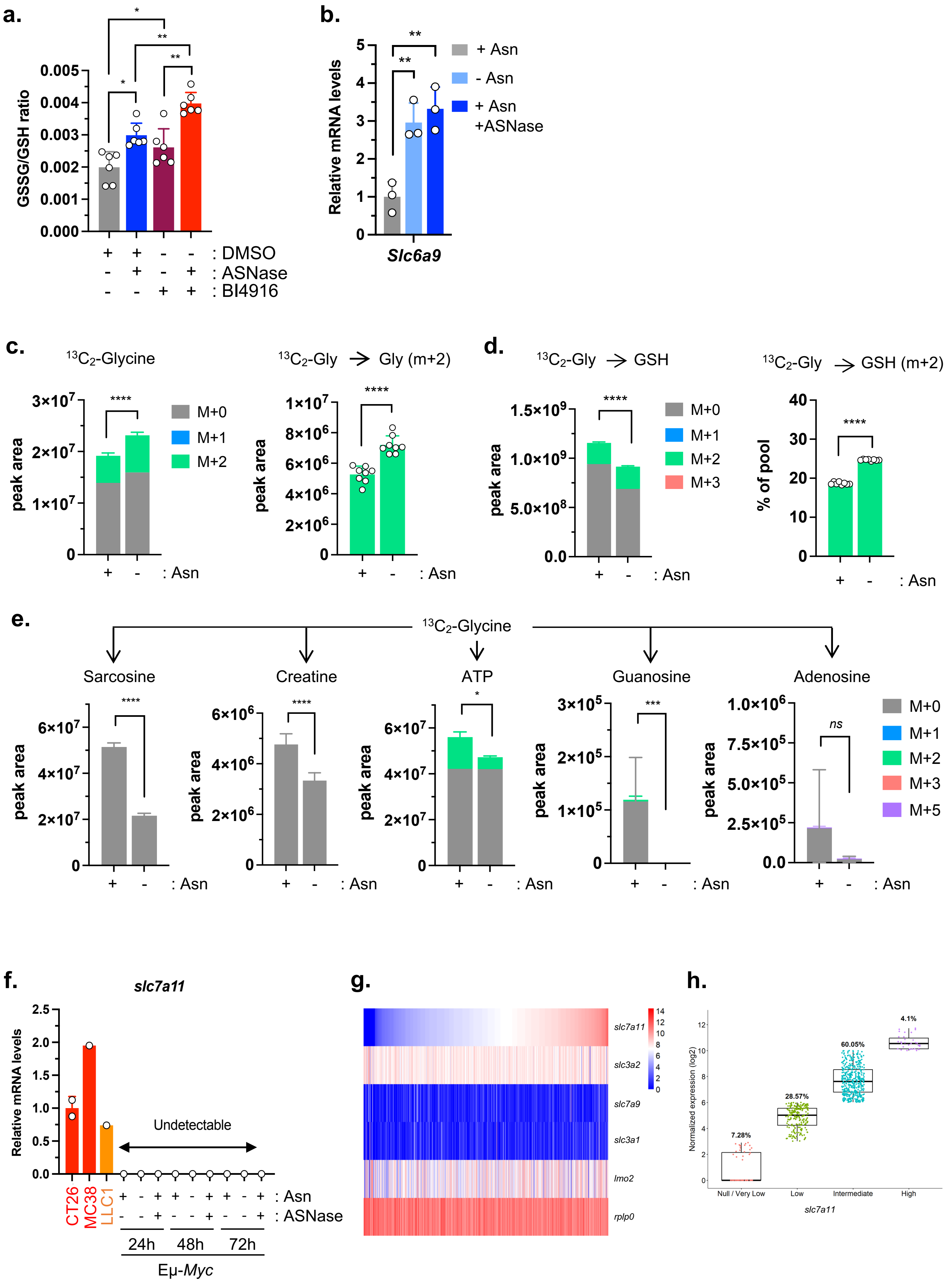


Aussel A. *et al.* Figure S6

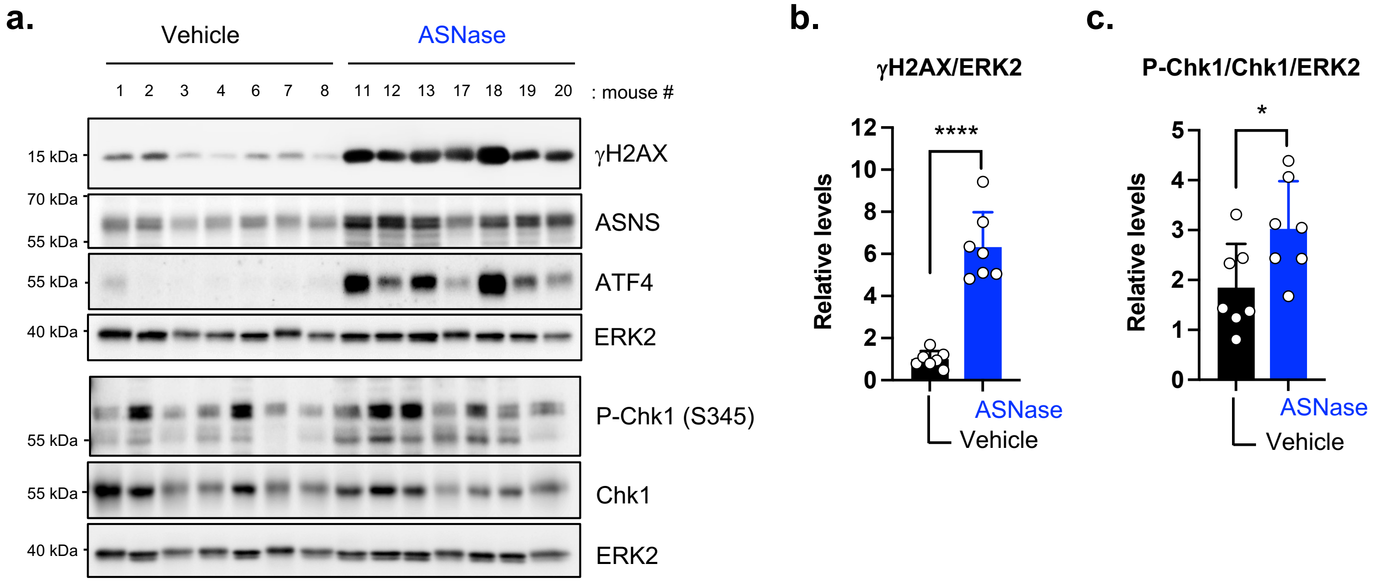


Aussel A. *et al.* Figure S7

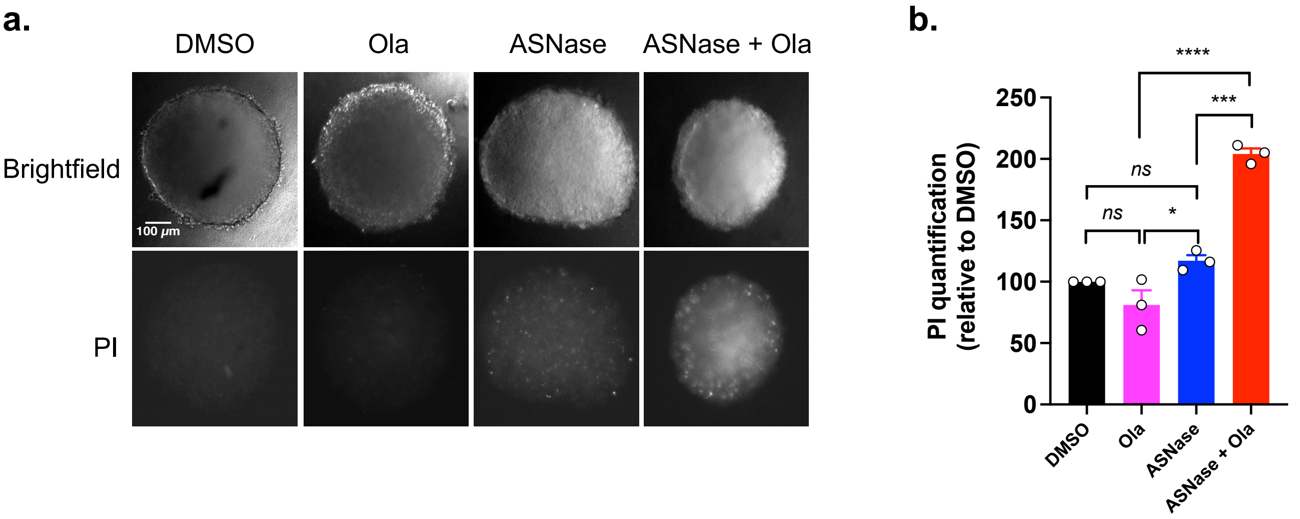


Aussel A. *et al.* Figure S8

**Table S1.** Statistically significant discriminant metabolites identified in B-cell lymphomas following *in vivo* treatment with Vehicle or ASNase, related to Figure 1.

**ASNase/Vehicle (from Eμ-*Myc* #506 cells-bearing mice)**

| | FC | log2(FC) | raw.pval | -log10(p) |
|---|---|---|---|---|
| Asparagine | 0.2205 | -2.1812 | 1.0825E-05 | 4.9656 |
| Glycine | 2.3121 | 1.2092 | 1.0825E-05 | 4.9656 |
| Serine | 1.9282 | 0.94722 | 1.0825E-05 | 4.9656 |
| UDP | 1.7214 | 0.78362 | 1.0825E-05 | 4.9656 |
| uridine | 0.60549 | -0.72382 | 1.0825E-05 | 4.9656 |
| Lactate | 0.68713 | -0.54134 | 1.0825E-05 | 4.9656 |
| CDP | 1.6816 | 0.74987 | 7.5776E-05 | 4.1205 |
| ADP | 1.4223 | 0.50819 | 7.5776E-05 | 4.1205 |
| GDP | 1.4468 | 0.53285 | 0.0001299 | 3.8864 |
| Sedoheptulose-7-phosphate (D) | 0.74055 | -0.43334 | 0.0001299 | 3.8864 |
| urate | 0.7928 | -0.33498 | 0.0001299 | 3.8864 |
| Glucose | 0.58394 | -0.7761 | 0.00020568 | 3.6868 |
| Eicosapentaenoic acid | 0.20275 | -2.3022 | 0.00032475 | 3.4884 |
| Tyrosine | 1.3512 | 0.43421 | 0.00032475 | 3.4884 |
| Glutamate | 0.76232 | -0.39152 | 0.00032475 | 3.4884 |
| Cystathionine | 1.8647 | 0.89892 | 0.00048713 | 3.3124 |
| Myristic acid | 0.54795 | -0.86788 | 0.00048713 | 3.3124 |
| SuccinylCysteine | 0.6834 | -0.54919 | 0.00048713 | 3.3124 |
| GTP | 2.814 | 1.4926 | 0.00072528 | 3.1395 |
| riboflavin | 0.72954 | -0.45494 | 0.00072528 | 3.1395 |
| beta-Alanine | 2.0161 | 1.0115 | 0.00105 | 2.9788 |
| G6P | 0.72947 | -0.45508 | 0.00105 | 2.9788 |
| N-carbamoyl-L-aspartic acid | 0.31784 | -1.6536 | 0.0015047 | 2.8226 |
| CTP | 2.1078 | 1.0757 | 0.0020892 | 2.68 |
| Lysine | 0.60158 | -0.73318 | 0.0020892 | 2.68 |
| Glycerol 3-phosphate | 0.62892 | -0.66906 | 0.0020892 | 2.68 |
| Glutamine | 0.66422 | -0.59027 | 0.0020892 | 2.68 |
| Pyruvate | 0.61995 | -0.68977 | 0.0028795 | 2.5407 |
| GSSG | 0.67143 | -0.57468 | 0.0028795 | 2.5407 |
| Arachidonic acid | 0.61227 | -0.70777 | 0.0038862 | 2.4105 |
| UTP | 1.4885 | 0.5739 | 0.0038862 | 2.4105 |
| UMP | 1.2647 | 0.33885 | 0.0038862 | 2.4105 |
| Adenosine | 1.5814 | 0.66124 | 0.005196 | 2.2843 |
| Decanoyl-Carnitine | 1.5425 | 0.62531 | 0.005196 | 2.2843 |
| ATP | 1.3973 | 0.48267 | 0.005196 | 2.2843 |
| Hexanoic acid | 0.72941 | -0.45521 | 0.005196 | 2.2843 |
| Butyryl-carnitine | 0.7747 | -0.3683 | 0.0068415 | 2.1649 |
| Butyric acid | 1.4527 | 0.53872 | 0.0089307 | 2.0491 |
| Acetyl CoA | 1.4165 | 0.50232 | 0.0089307 | 2.0491 |
| Palmitoleic acid | 0.64387 | -0.63515 | 0.011496 | 1.9394 |
| cis-aconitate | 0.76917 | -0.37863 | 0.011496 | 1.9394 |
| Octanoyl-carnitine | 1.4907 | 0.576 | 0.01469 | 1.833 |
| Ribose | 0.74609 | -0.42258 | 0.01469 | 1.833 |
| Ornithine | 0.71762 | -0.4787 | 0.018543 | 1.7318 |
| Methylhistidine | 1.5334 | 0.61669 | 0.023231 | 1.6339 |
| Xanthosine | 0.7131 | -0.48782 | 0.023231 | 1.6339 |
| acetyl-carnitine | 0.74095 | -0.43255 | 0.023231 | 1.6339 |
| guanosine | 1.2686 | 0.34323 | 0.023231 | 1.6339 |
| 5-Hydroxylysine | 0.45443 | -1.1379 | 0.028806 | 1.5405 |
| Fructose | 1.2779 | 0.35382 | 0.028806 | 1.5405 |
| Palmitic acid | 0.68522 | -0.54537 | 0.035463 | 1.4502 |
| Succinyladenosine | 2.2705 | 1.183 | 0.043257 | 1.3639 |

**ASNase vs Vehicle (from Eμ-*Myc* #688 cells-bearing mice)**

| | FC | log2(FC) | raw.pval | -log10(p) |
|---|---|---|---|---|
| Asparagine | 0.14412 | -2.7947 | 7.396E-07 | 6.131 |
| Glycine | 2.2911 | 1.196 | 5.1772E-06 | 5.2859 |
| Serine | 2.0535 | 1.0381 | 5.1772E-06 | 5.2859 |
| Threonine | 1.7526 | 0.80949 | 2.2188E-05 | 4.6539 |
| CDP | 1.9014 | 0.92704 | 3.3282E-05 | 4.4778 |
| GDP | 2.0174 | 1.0125 | 4.9553E-05 | 4.3049 |
| L-Alanine | 1.8877 | 0.91666 | 4.9553E-05 | 4.3049 |
| L-Sarcosine | 1.8877 | 0.91666 | 4.9553E-05 | 4.3049 |
| Ornithine | 1.8255 | 0.8683 | 4.9553E-05 | 4.3049 |
| Proline | 1.6086 | 0.68584 | 4.9553E-05 | 4.3049 |
| Quinolinic acid | 0.47647 | -1.0695 | 7.1741E-05 | 4.1442 |
| O-Phosphoethanolamine | 1.3628 | 0.44653 | 0.0001028 | 3.988 |
| 3-hydroxybutyric acid | 0.19027 | -2.3939 | 0.00027439 | 3.5616 |
| Phenylalanine | 1.489 | 0.57435 | 0.00027439 | 3.5616 |
| IMP | 1.4959 | 0.58106 | 0.00053052 | 3.2753 |
| acetyl-carnitine | 0.68793 | -0.53966 | 0.00085794 | 3.0665 |
| beta-Alanine | 2.1206 | 1.0845 | 0.00089861 | 3.0464 |
| Inosine | 1.9795 | 0.98516 | 0.0011146 | 2.9529 |
| Hexanoyl-carnitine | 0.54452 | -0.87694 | 0.0011146 | 2.9529 |
| aKG | 0.59446 | -0.75036 | 0.0011146 | 2.9529 |
| CMP | 1.2716 | 0.34662 | 0.0011146 | 2.9529 |
| Octanoyl-carnitine | 0.50281 | -0.99191 | 0.0014326 | 2.8439 |
| Dodecanoyl-carnitine | 0.52919 | -0.91814 | 0.0018298 | 2.7376 |
| Decanoyl-Carnitine | 0.5271 | -0.92384 | 0.0023164 | 2.6352 |
| Docosahexaenoic acid | 1.7242 | 0.78591 | 0.002914 | 2.5355 |
| Valine | 1.6477 | 0.72046 | 0.002914 | 2.5355 |
| guanosine | 1.5824 | 0.66213 | 0.002914 | 2.5355 |
| Pyruvate | 0.70938 | -0.49537 | 0.002914 | 2.5355 |
| Argininosuccinate | 1.924 | 0.94411 | 0.0032282 | 2.491 |
| Creatinine | 1.2787 | 0.35468 | 0.0045131 | 2.3455 |
| ADP | 1.3023 | 0.38107 | 0.0045799 | 2.3391 |
| hypoxanthine | 1.4403 | 0.52636 | 0.0055596 | 2.255 |
| Adenine | 1.3678 | 0.45187 | 0.0055596 | 2.255 |
| Aspartate | 1.3567 | 0.44012 | 0.0055596 | 2.255 |
| 5-oxo-L-Proline | 1.3996 | 0.485 | 0.0055733 | 2.2539 |
| L-Dihydroorotic acid | 0.3005 | -1.7345 | 0.0082932 | 2.0813 |
| cis-aconitate | 0.73013 | -0.45377 | 0.0082932 | 2.0813 |
| Myristoyl-carnitine | 0.63681 | -0.65106 | 0.012094 | 1.9174 |
| Propionyl-carnitine | 1.3643 | 0.44818 | 0.012094 | 1.9174 |
| Aminoadipate | 1.3412 | 0.42356 | 0.012094 | 1.9174 |
| Leucine | 1.4162 | 0.50201 | 0.014493 | 1.8389 |
| UDP | 1.4147 | 0.50049 | 0.014493 | 1.8389 |
| Succinic acid | 0.69876 | -0.51712 | 0.015292 | 1.8155 |
| Cystathionine | 2.0429 | 1.0306 | 0.020489 | 1.6885 |
| Butyric acid | 1.4997 | 0.58471 | 0.020489 | 1.6885 |
| Adenosine | 1.3644 | 0.44826 | 0.020489 | 1.6885 |
| ATP | 1.2544 | 0.32695 | 0.024184 | 1.6165 |
| N-carbamoyl-L-aspartic acid | 0.43636 | -1.1964 | 0.028421 | 1.5464 |
| GTP | 1.9906 | 0.99322 | 0.028421 | 1.5464 |
| Pantothenate | 1.4903 | 0.57565 | 0.028421 | 1.5464 |
| Tyrosine | 1.298 | 0.37632 | 0.028421 | 1.5464 |
| Tryptophan | 1.2501 | 0.32207 | 0.028421 | 1.5464 |
| nicotinamide N-oxide | 1.4419 | 0.52794 | 0.033241 | 1.4783 |
| S-Adenosyl-L-methionine | 1.2516 | 0.32382 | 0.033241 | 1.4783 |
| Palmitoyl-carnitine | 0.65787 | -0.60414 | 0.038721 | 1.4121 |
| FAD | 1.3347 | 0.41653 | 0.038721 | 1.4121 |
| Oleic acid | 0.66373 | -0.59134 | 0.044902 | 1.3477 |
| Methionine | 1.3419 | 0.42423 | 0.044902 | 1.3477 |

**Table S2.** Fold changes in proteinogenic amino acids levels in B-cell lymphomas treated *in vivo* with ASNase *vs* Vehicle, related to Figure 1.

**ASNase/Vehicle (from Eµ-*Myc* #506 cells-bearing mice)**

| | | | Fold Change | log2(FC) | p.value | -log10(p) |
|---|---|---|---|---|---|---|
| NEAA | Down | **Asparagine** | **0.2205** | **-2.1812** | **1.08E-05** | **4.9656** |
| | | **Glutamine** | **0.66422** | **-0.59027** | **0.0020892** | **2.68** |
| | | **Glutamate** | **0.76232** | **-0.39152** | **0.00032475** | **3.4884** |
| | Up | **Tyrosine** | **1.3512** | **0.43421** | **0.00032475** | **3.4884** |
| | | **Serine** | **1.9282** | **0.94722** | **1.08E-05** | **4.9656** |
| | | **Glycine** | **2.3121** | **1.2092** | **1.08E-05** | **4.9656** |
| | Non significant | Aspartate | 0.77865 | -0.36095 | 0.075256 | 1.1235 |
| | | Arginine | 0.88282 | -0.17981 | 0.24745 | 0.60651 |
| | | Proline | 1.1041 | 0.1429 | 0.16549 | 0.78122 |
| | | Alanine | 1.2225 | 0.2898 | 0.063013 | 1.2006 |
| EAA | Down | **Lysine** | **0.60158** | **-0.73318** | **0.0020892** | **2.68** |
| | | **Methionine** | **0.80007** | **-0.3218** | **0.005196** | **2.2843** |
| | | **IsoLeucine** | **0.84453** | **-0.24377** | **0.043257** | **1.3639** |
| | Up | **Threonine** | **1.2303** | **0.29903** | **0.01469** | **1.833** |
| | Non significant | Leucine | 0.88994 | -0.16822 | 0.10512 | 0.9783 |
| | | Valine | 0.91508 | -0.12802 | 0.12301 | 0.91008 |
| | | Histidine | 0.93239 | -0.10099 | 0.39305 | 0.40555 |
| | | Tryptophan | 0.98528 | -0.021389 | 0.57874 | 0.23752 |
| | | Phenylalanine | 1.0517 | 0.072731 | 0.21756 | 0.66242 |

**ASNase/Vehicle (from Eµ-*Myc* #688 cells-bearing mice)**

| | | | Fold Change | log2(FC) | p.value | -log10(p) |
|---|---|---|---|---|---|---|
| NEAA | Down | **Asparagine** | **0.14412** | **-2.7947** | **7.40E-07** | **6.131** |
| | Up | **Tyrosine** | **1.298** | **0.37632** | **0.028421** | **1.5464** |
| | | **Aspartate** | **1.3567** | **0.44012** | **0.0055596** | **2.255** |
| | | **Proline** | **1.6086** | **0.68584** | **4.96E-05** | **4.3049** |
| | | **Alanine** | **1.8877** | **0.91666** | **4.96E-05** | **4.3049** |
| | | **Serine** | **2.0535** | **1.0381** | **5.18E-06** | **5.2859** |
| | | **Glycine** | **2.2911** | **1.196** | **5.18E-06** | **5.2859** |
| | Non significant | Glutamate | 1.0351 | 0.049833 | 0.37769 | 0.42287 |
| | | Glutamine | 1.1569 | 0.21031 | 0.29134 | 0.5356 |
| | | Arginine | 1.2063 | 0.27054 | 0.12769 | 0.89385 |
| EAA | Up | **Histidine** | **1.1879** | **0.24837** | **0.014493** | **1.8389** |
| | | **Tryptophan** | **1.2501** | **0.32207** | **0.028421** | **1.5464** |
| | | **Methionine** | **1.3419** | **0.42423** | **0.044902** | **1.3477** |
| | | **Leucine** | **1.4162** | **0.50201** | **0.014493** | **1.8389** |
| | | **Phenylalanine** | **1.489** | **0.57435** | **0.00027439** | **3.5616** |
| | | **Valine** | **1.6477** | **0.72046** | **0.002914** | **2.5355** |
| | | **Threonine** | **1.7526** | **0.80949** | **2.22E-05** | **4.6539** |
| | Non significant | Lysine | 1.1024 | 0.14067 | 0.41882 | 0.37797 |
| | | IsoLeucine | 1.267 | 0.34143 | 0.12769 | 0.89385 |
